# Transport mechanism of DgoT, a bacterial homolog of SLC17 organic anion transporters

**DOI:** 10.1101/2024.02.07.579339

**Authors:** Nataliia Dmitrieva, Samira Gholami, Claudia Alleva, Paolo Carloni, Mercedes Alfonso-Prieto, Christoph Fahlke

## Abstract

The solute carrier 17 (SLC17) family contains anion transporters that accumulate neuro-transmitters in secretory vesicles, remove carboxylated monosaccharides from lysosomes, or extrude organic anions from the kidneys and the liver. We combined classical molecular dynamics simulations, Markov state modeling and hybrid first principles quantum mechani-cal/classical mechanical (QM/MM) simulations with experimental approaches to describe the transport mechanisms of a model bacterial protein, the D-galactonate transporter DgoT, at atomic resolution. We found that protonation of D46 and E133 precedes galactonate binding and that substrate binding induces closure of the extracellular gate, with the conserved R47 coupling substrate binding to transmembrane helix movement. After isomerization to an inward-facing conformation, deprotonation of E133 and subsequent proton transfer from D46 to E133 opens the intracellular gate and permits galactonate dissociation either in its unprotonated form or after proton transfer from E133. After release of the second proton, apo DgoT returns to the outward-facing conformation. Our results provide a framework to understand how various SLC17 transport functions with distinct transport stoichiometries can be attained through subtle variations in proton and substrate binding/unbinding.

## Introduction

Solute carrier 17 (SLC17) transporters fulfil a variety of cellular functions (Reimer 2013; Omote et al. 2016; Li et al. 2022). They transport diverse anionic substrates; mostly electro-chemical proton (H^+^) gradients as the driving force, with a large variability in transport stoi-chiometry ranging from electrogenic H^+^-glutamate exchange by VGLUT (Kolen et al. 2023) to electroneutral H^+^-sialic acid symport by sialin (Morin et al. 2004; Hu et al. 2023). Although this protein family has been studied for several decades (Kaback and Guan 2019), the mechanistic basis of SLC17 organic anion transport remains insufficiently understood.

The bacterial homolog DgoT (from *E. coli*) transports negatively charged galactonate in symport with more than one proton and, thus, differs from mammalian SLC17s in both its main substrate and transport stoichiometry (Leano et al. 2019). It belongs to the large major facilitator superfamily (MFS). MFS family members exhibit a distinct topology that includes 12 transmembrane (TM) helices, with the substrate-binding pocket located between two pseudo-symmetrical 6-TM bundles in the N-domain (TM1–TM6) and C-domain (TM7– TM12) (Quistgaard et al. 2016; Drew et al. 2021) . DgoT has been crystallized in two distinct conformations: (i) inward-facing and (ii) outward-facing with galactonate bound to the central binding site (Leano et al. 2019). These structures revealed four charged residues within the transmembrane domains, D46, R47, R126, and E133, with the last three conserved across the SLC17 family. The high structural similarity with vesicular glutamate transporters (VGLUTs) (Li et al. 2020) and lysosomal sialic acid transporter (Hu et al. 2023) (Supplementary Figs 1 and 2), together with differences in substrate selectivity and transport stoichiometry, makes DgoT an attractive model to study the atomistic basis of coupled transport in the SLC17 family. Here, we present a comprehensive study of DgoT transport using a combination of *in vitro* experiments and molecular dynamics (MD) simulations.

**Figure 1.**
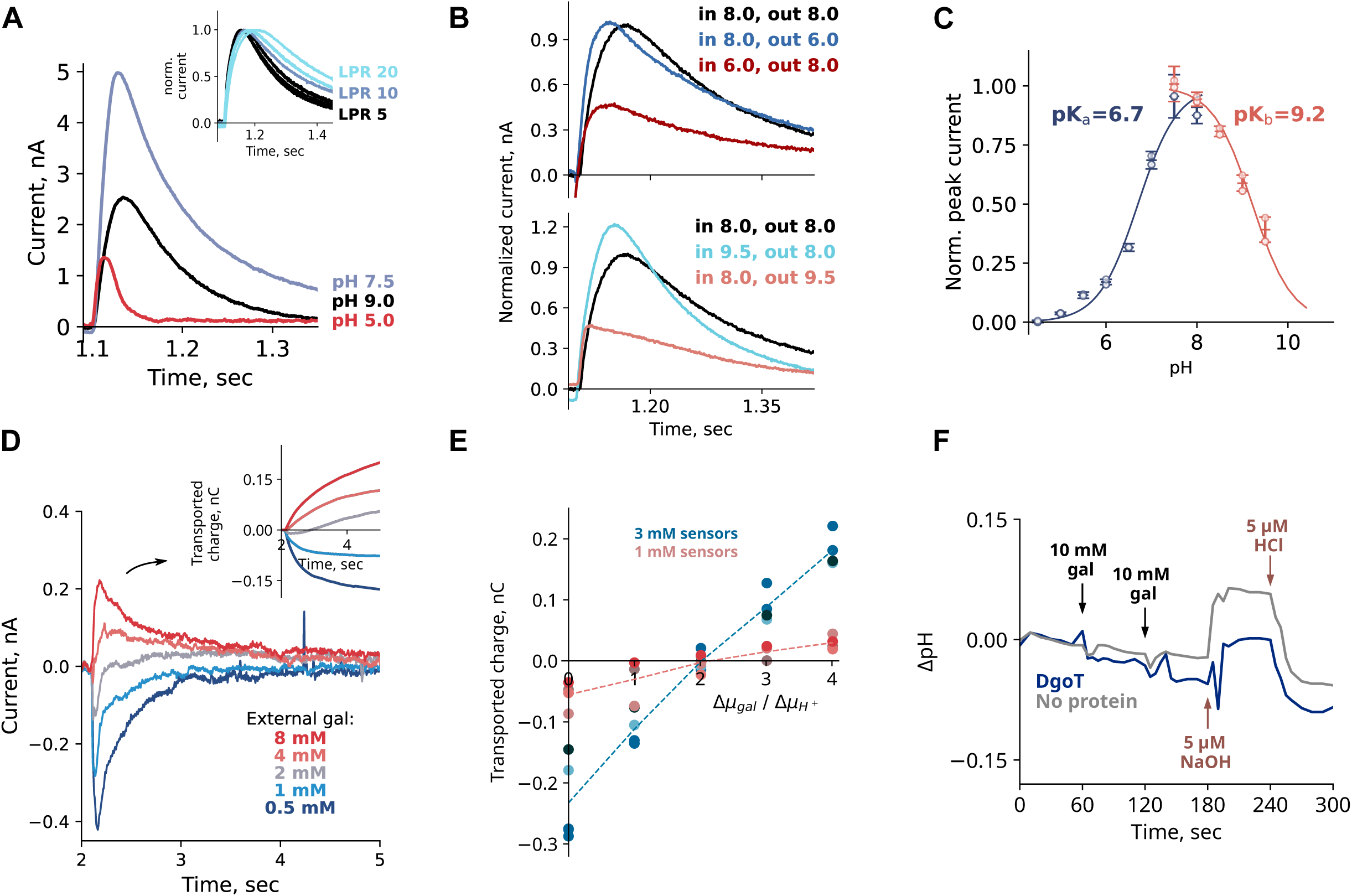
Functional characterization of DgoT. A pH dependence of WT DgoT peak currents measured by SSME upon application of 10 mM D-galactonate concentration jump. The inset shows transport currents at pH 8.0 from liposomes reconstituted with different lipid to protein ratios (LPR). Currents were normalized to their peak value for comparison of the current decay. B Transient currents under symmetrical (black line) or asymmetrical conditions. Peak current decreases if acidic pH is applied inside liposomes (top) or alkaline pH is applied outside (bottom). C Peak currents and determination of apparent pK values. The solid lines are fits to the data using the equations described in Methods. Experiments were performed in duplicate on two independent sensors. D Representative transport current traces used for the determination of the transport stoichiometry. The inset shows time dependence of transported charge obtained by integration of current traces. E Plots of transported charge *versus* ratio of the galactonate and proton chemical potentials. F Changes in pH after addition of substrate to purified DgoT in detergent. Additions of 10 mM of galactonate are indicated by the black arrows. 5 µM HCl or NaOH were added to induce a pH shift comparable with the expected changes in H^+^ concentration due to proton binding to DgoT.

**Figure 2.**
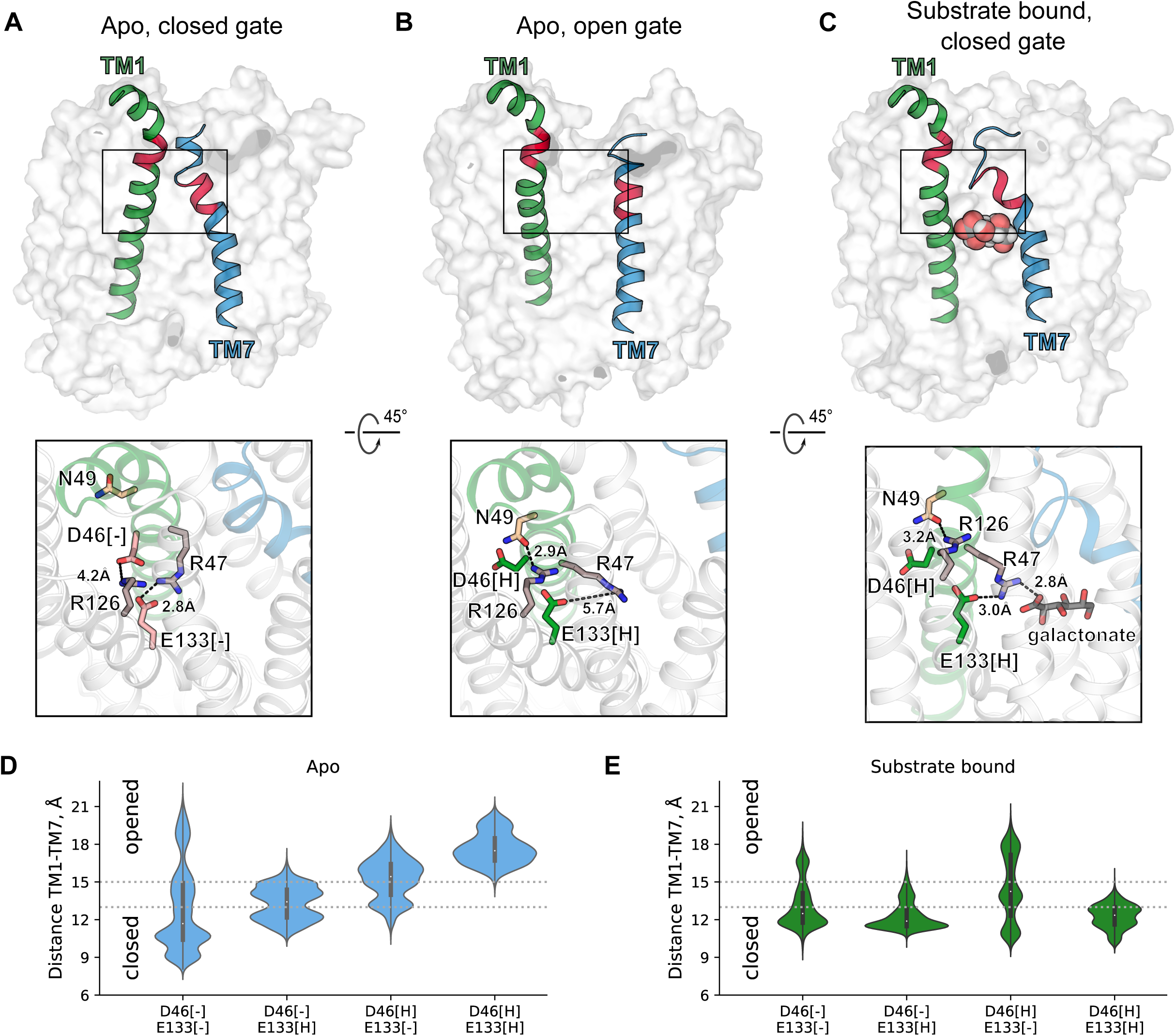
Unbiased MD simulations reveal protonation- and substrate-dependent dynamics of extracellular gate. A-C Top row: snapshots showing the arrangement of the gating helices TM1 and TM7 in unbiased MD simulations with DgoT in outward-facing conformation with D46 and E133 deprotonated (A), protonated (B) and galactonate bound (C). The red colored parts of the helices correspond to residues 48-52 (TM1) and residues 271-275 (TM7). Bottom row: representative snapshots showing interactions around D46 and E133. D, E Probability densities for extracellular gate opening in *apo* (D) and galactonate-bound (E) MD simulations with DgoT in the outward-facing conformation with different protonation states of D46 and E133. The TM1-TM7 distance is measured as distance between center of mass of Ca atoms of residues 48-52 and residues 271-275.

## Results

### DgoT mediates the coupled symport of two H^+^ and one D-galactonate

We studied DgoT transport using solid-supported membrane-based electrophysiology (SSME) with purified DgoT reconstituted into proteoliposomes. In such recordings, transport activity is induced by fast solution exchange. The resulting changes in proteoliposomal membrane potential are converted into an electrical signal and detected by a measuring electrode via capacitive coupling (Schulz et al. 2008; Bazzone et al. 2021).

Fig. 1a depicts SSME currents after galactonate concentration jumps at three different pHs. At neutral and alkaline pH, the time course for current decay decreased at higher lipid–protein ratios (LPRs; illustrated for pH 8.0 in the inset). This result indicates that the measured currents are caused by electrogenic DgoT transport (Bazzone et al. 2021). Transport currents assumed a maximum amplitude at pH 7.5 (blue line) and decreased at higher pH values (Fig. 1a; pH 9.0, black line). Under asymmetrical ionic conditions (i.e., with an alkaline pH outside, but not inside, the liposomes), the peak current was reduced (Fig. 1b bottom). This suggests that reduced transport at alkaline pH is caused by proton depletion at the binding site outside the liposomes. Under symmetrical pH conditions, the current decay was biphasic at pH values lower than 7.5 (Supplementary Fig. 3): the fast component represents conformational changes triggered by galactonate binding and the slow component is associated with the transport activity of the protein. At pH values lower than 6, only the pre-steady state, but not the transport component, was observed (Fig. 1a; red line). Acidic pH inside, but not outside, the liposomes inhibits transport activity, as demonstrated under asymmetrical ionic conditions (Fig. 1b top). We conclude that current reduction under conditions of acidic or alkaline pH is regulated by two different processes: proton binding from the outside and proton release to the inside of the proteoliposomes, respectively. Thus, analysis of the peak currents allowed the determination of apparent pKa values for proton binding and release (Fig. 1c).

The transport currents generated by DgoT were positive (Fig. 1a), indicating that at least two protons are transported with each galactonate molecule, as previously reported (Leano et al. 2019). We used a reversal potential assay to determine the transport stoichiometry (Thomas et al. 2021). In SSME, no voltage is applied to the membrane and concentration gradients are the only driving force for coupled transport. The free energy for the coupled transport reaction is given by:

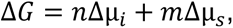

where *n* and *m* are number of protons and galactonate molecules transported together. The current reverses at Δ*G* = 0:

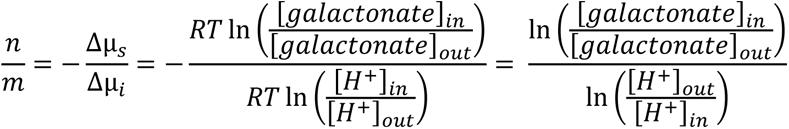

Thus, plotting the transported charge against the ratio of the electrochemical gradients enables the transport stoichiometry to be determined (Thomas et al. 2021).

We preloaded proteoliposomes with galactonate-containing internal solutions and applied sets of external solutions to generate an outwardly directed proton concentration gradient at various galactonate concentration gradients. The currents obtained with empty liposomes—used as the negative control (Supplementary Fig. 4)—were subtracted. Null transport occurred at the gradient ratios corresponding to a stoichiometry of 2 H^+^:1 galactonate (Fig. 1d, e).

To investigate the binding order of H^+^ and galactonate, we measured changes to the external pH of solutions containing purified detergent-solubilized protein upon the addition of galactonate (Soskine et al. 2004). If galactonate binding precedes proton association, we would expect the pH to increase upon galactonate addition. However, addition of 10 mM galactonate to purified DgoT in unbuffered detergent-containing solutions did not elicit any change in pH (Fig. 1f). This result suggests that the substrate binds to the protonated trans-porter and does not induce protonation of DgoT. Therefore, we conclude that DgoT binds two H^+^ prior to galactonate association.

### Protonation of key residues regulates extracellular gate dynamics

We studied H^+^ and galactonate binding to outward-facing DgoT using unbiased all-atom MD simulations. For this, we used the crystal structure of E133Q DgoT (Leano et al. 2019) (PDB ID: 6E9O) as the starting conformation after reverting the mutation. First, we studied the effect of protonation on *apo* protein dynamics. The crystallographic galactonate molecule was removed from the binding site, and four different systems were generated with D46 and E133 individually protonated or deprotonated. We observed reproducible movements of the TM1 and TM7 transmembrane helices, with most flexibility in the most extracellular segments (around residues 48–52 of TM1 and 271–275 of TM7). This behavior resembles the proposed role of TM1 and TM7 as gating helices that mediate transitions between open and occluded conformations, as in other MFS members (Smirnova et al. 2014; Qureshi et al. 2020; Feng et al. 2021).

When D46 and E133 were unprotonated, the extracellular gate (defined as distance between the center of mass of the Cα atoms of residues 48–52 and residues 271–275) was mostly closed (Fig. 2a, d). Protonation of both D46 and E133 locked the gate in an open conformation (Fig. 2b, d), and protonation of only one of these residues only partially opened the gate (Fig. 2d). These changes in position of the gating helices were coupled to local rearrangements in the N-terminal domain. Charged, but not protonated, D46 and E133 interact with R126 and R47, respectively. After protonation, the side chain of D46 moves away, allowing R126 to interact with N49. This results in rotation of TM1 and stabilization of the open state of the extracellular gate (compare Fig. 2a, b). In addition, protonation of E133 releases R47, which then becomes available to interact with the carboxyl group of galactonate.

We performed unbiased simulations with the *apo* outward-facing structure in presence of 100 mM galactonate in solution. Before starting the MD simulations, D46 and E133 were protonated or deprotonated individually. For each system, at least one of the two key residues was protonated (five replicates were used). We observed spontaneous galactonate-binding events in two simulations in which both D46 and E133 were protonated and in one simulation in which only E133 was protonated. In all cases, opening of the extracellular gate preceded the entry of galactonate into the binding site (Fig. 3d).

**Figure 3.**
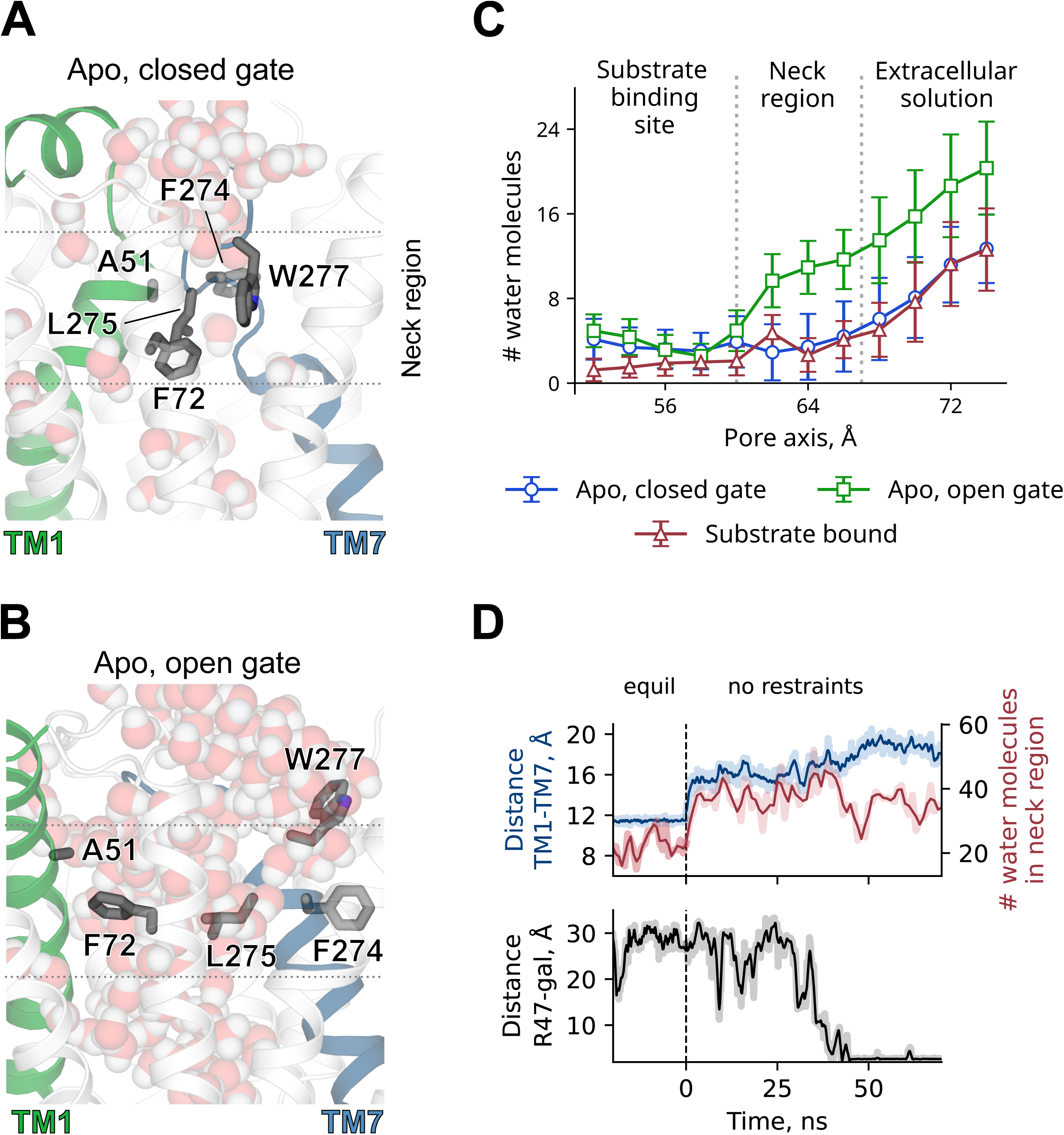
Hydrophobic interactions at the level of the extracellular gate. A, B Snapshots showing water molecules distribution in simulations with DgoT in outward-facing conformation with closed (A) and open (B) extracellular gate. Representative snapshots were taken from unbiased MD simulations with both D46 and E133 either deprotonated (A) or protonated (B). C Hydration profile of the pore represented by the number of water molecules in 2 Å sections along the pore axis. Trajectories with following parameters were used for analysis: *apo*, closed gate – D46 and E133 protonated; *apo*, open gate and substrate bound – D46 and E133 protonated. In *apo* simulations only frames with minimum distance between side chains of F72 and W277 < 12.5 Å (blue line) or > 12.5 Å (green line) were used. D Time course of extracellular gate opening (measured as TM1-TM7 distance, as in Fig. 2), number of water molecules in 10 Å section near extracellular gate (z coordinate between 58 and 68 Å) and galactonate binding to DgoT (measured as minimum distance between galac-tonate molecule and guanidinium group of R47) in a trajectory with both D46 and E133 protonated. Shaded lines represent raw data from the trajectory, solid lines are moving averages.

### Substrate binding induces closure of the extracellular gate

Unbiased simulations of outward-facing DgoT with bound galactonate revealed the reduced flexibility of gating helices TM1 and TM7 (Fig. 2c) and the extracellular gate is more likely to be in a closed conformation (Fig. 2e). Although the gate could close regardless of the protonation state of key residues in substrate-bound DgoT, the equilibrium was shifted toward the conformation with closed extracellular gate in simulations with only E133 protonated or with both D46 and E133 protonated (Fig. 2e).

Protonation of E133 permits the R47 side chain to simultaneously interact with the carboxyl groups of galactonate and E133, resulting in a more compact arrangement of the charged residues (Fig. 2c). Substrate interaction with residues on both gating helices facilitates bending of the TM7 extracellular segment toward TM1. Hydrophobic interactions also play a role in forming occluded conformations. In the outward-facing occluded state, the substrate-binding site is separated from the extracellular solution by hydrophobic residues located in TM1, TM2, and TM7 (F72, F274, and W277, respectively; Fig. 3a). When the extracellular gate opens, the position of the side chains of the three hydrophobic residues changes to allow more water molecules to flow between TM1 and TM7 (Fig. 3b) and galactonate to enter the binding site (Fig. 3c). This resembles a “clamp-and-switch” mechanism (Quistgaard et al. 2016), in which occlusion of the binding site precedes a rocker-switch-type rotation on the N- and C-terminal domains, causing the binding site to become exposed to the opposite side (Qureshi et al. 2020).

### Structural basis of substrate release

We next ran unbiased simulations with the inward-facing structure of DgoT after placing a galactonate molecule into the binding site. The position for the substrate molecule was determined by aligning the crystal structures of the *apo* protein in the inward-facing state (PDB ID: 6E9N) with the galactonate-bound outward-facing state (PDB ID: 6E9O). Galactonate in solution is predicted to have a pKa of 3.9 and, therefore, should bind to outward-facing DgoT in its deprotonated form. However, we also considered the possibility that galactonate could be protonated in the binding pocket due to changes in the local environment and then released in its protonated form. In 7 out of 38 individual 500 ns unbiased MD simulations, we observed spontaneous galactonate release (Supplementary Table 1). In all cases, the substrate passed between TM4 and TM10, which act as gating helices on the intracellular side of the protein. The intracellular segments of these helices (around residues 139–143 and 372–375, respectively) showed a high degree of flexibility in these simulations. The side chains of F137 and W373 formed a hydrophobic lock similar to the one at the extracellular gate (Fig. 4a), and galactonate could only pass through this neck region if a water-filled pore was formed between the side chains of F137 and W373 (Fig. 4b, c).

**Figure 4.**
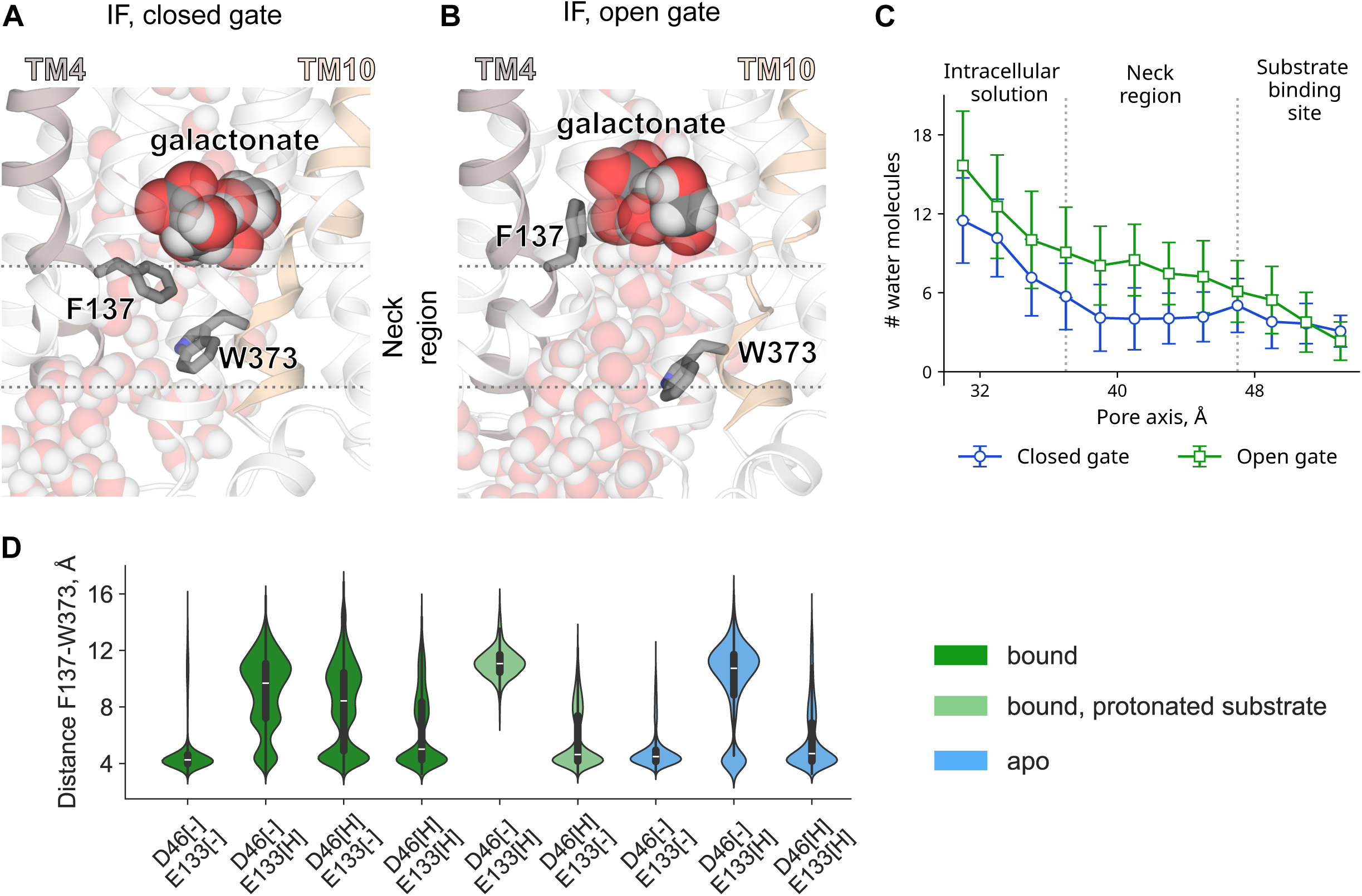
A hydrophobic lock within the intracellular gate regulates access to the binding site. A, B Snapshots showing water molecule distribution in simulations with DgoT in inward-facing conformation with closed (A) and open (B) intracellular gate. Representative snapshots were taken from unbiased MD simulations with D46 and E133 protonated (A) or only E133 protonated (B). C Hydration profile of the pore, represented by the number of water molecules in 2-Å sections along the pore axis. Trajectories with following parameters were used for analysis: closed gate – D46 and E133 protonated; open gate – D46 deprotonated, E133 protonated. Substrate was bound to the protein in both cases. Only frames with minimum distance between side chains of F137 and W373 < 8 Å (blue line) or > 8 Å (green line) were used. D Probability densities for intracellular gate opening (measured as distance between side chains of F137 and W373) in MD simulations with DgoT in inward-facing conformation with different protonation states of D46 and E133 and galactonate and various substrate occupancies.

All spontaneous release events were observed in systems containing deprotonated D46, which suggests that the proton must be released from this residue prior to galactonate unbinding. Fig. 4d shows probability densities for intracellular gate opening (measured as the minimum distance between the side chains of F137 and W373) for various protonation and substrate-binding states. The intracellular gate mostly assumed a closed conformation in the substrate-bound double protonated system. Deprotonation of D46 favored the open state of the gate in systems with a protonated or deprotonated substrate. When galactonate is protonated, the intracellular gate has limited flexibility and becomes locked in the open state, when only E133 is protonated, and in the closed state, when only D46 is protonated. After release of the substrate and both H^+^ ions, the intracellular gate closes, permitting reorientation of the *apo* transporter to the outward conformation. We conclude that D46 deprotonation increases the probability of galactonate release from the inward-facing DgoT by promoting the opening of the intracellular gate.

### Proton release from inward-facing DgoT

Galactonate spontaneously dissociated from inward-facing DgoT only when D46 was deprotonated (Supplementary Table 1). However, in MD simulations of DgoT in which both D46 and E133 were protonated, the D46 carboxyl group was sequestered, making direct deprotonation to the intracellular solution unlikely (Fig. 5a). In addition, the electrostatic potential (calculated using *g_elpot* (Kostritskii et al. 2021; Kostritskii and Machtens 2023)) was higher on the E133 carboxyl group than on D46 (Fig. 5b), indicating that E133 is likely the first residue to release its proton. These observations suggest D46 donates its proton to E133 after the latter had released its proton.

**Figure 5.**
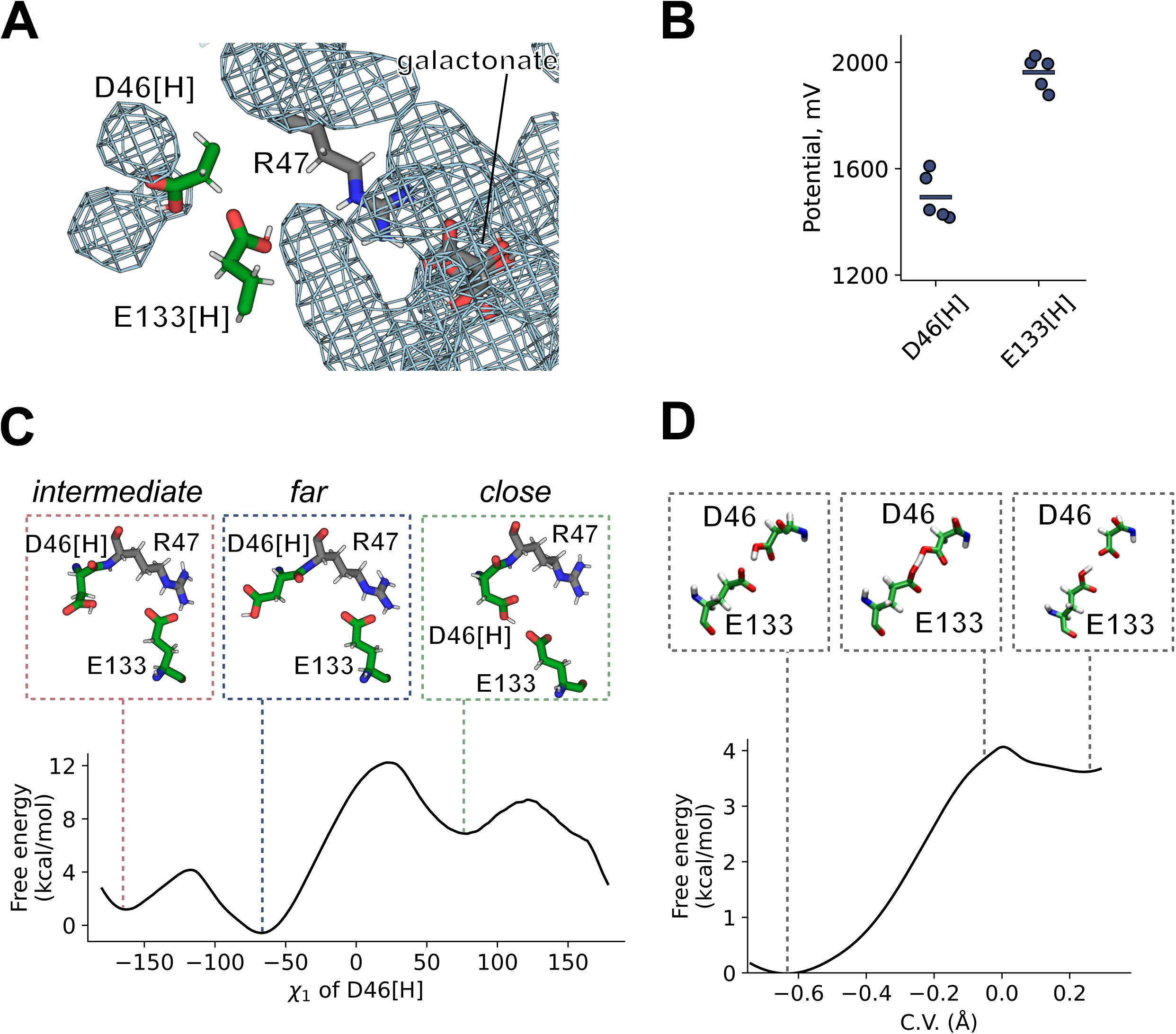
Proton release from D46. A Water occupancy map in unbiased simulations with D46 and E133 protonated, contoured at an occupancy level of 0.1. B Electrostatic potential at the carboxyl groups of protonated D46 and E133 in the unbiased MD simulation. C Classical MTD simulations of the conformational change of protonated D46. D Free energy profile for the proton transfer between D46 and E133, computed at the QM (BLYP)/MM level. The insets show representative starting, transition state, and final configurations. Error bars are omitted since they are smaller than marker size

Proton transfer from D46 requires a conformational change of its side chain, so as to be in close proximity to E133; therefore, we calculated the free energy associated with conformational change as a function of the N-Cα-Cβ*-Cγ* (χ_1_) dihedral angle of D46 using well-tempered metadynamics (WT-MTD) (Tribello et al. 2014). Fig. 5c shows the presence of a minimum at χ_1_ = -70°, corresponding to the conformation sampled in unbiased MD simulations. However, another minimum is present (hereafter denoted as *close*-D46, characterized by χ_1_ = +70°), which has a higher free energy (by ∼8 kcal mol^−1^) and is separated by a ∼12 kcal mol^−1^ barrier. In this conformation, the carboxyl group of D46 is H-bonded to E133, enabling proton transfer to take place.

> E133 may release its proton to either the solvent or galactonate (Fig. 5a). Hydrated excess protons can reorganize the surrounding water wires and facilitate further proton translocation (Peng et al. 2015; Li and Voth 2021); therefore, we first studied proton transfer from E133 to the adjacent water molecule to form a hydronium ion. Quantum mechanics/molecular mechanics (QM/MM) thermodynamics integration (TI) calculations showed that the free energy barrier can be as small as 8 kcal mol^−1^ (Supplementary Fig. 5). Moreover, in the presence of the ion pair between the negatively charged E133 and the adjacent hydronium ion, the surrounding water molecules connect the excess proton with both galactonate and the solution. To represent the diverse range of possible H-bond networks, we selected seven structures (see Methods) for use in unbiased QM/MM MD simulations in which the quantum part was treated at the density functional theory (DFT)-BLYP level (Becke 1988). In simulations with three of the selected structures, the galactonate carboxyl group accepted the proton after a few picoseconds, either via water molecules only or via water molecules along with the substrate hydroxyl group (Supplementary Figure 6 and Supplementary Table 2). In simulations with another three structures, the proton was transferred to one of the molecules of the water wire, without substrate protonation. In simulations with the seventh structure, the proton was shared between the substrate and a neighboring water molecule. Therefore, despite the limited statistics and underestimation of the proton transfer barriers due to the BLYP functional (Mangiatordi et al. 2012), we suggest that proton transfer to galactonate or the solvent is viable over a sub-nanosecond timescale. The released proton might eventually leave the protein either bound to the substrate or through the solvent via a Grotthuss mechanism.Finally, we performed QM/MM TI calculations to study the energetics associated with a proton transfer from *close*-D46 to E133. There was a small free energy barrier (around 4 kcal mol^−1^) associated with D46 deprotonation (Fig. 5d). Given the overall moderate energy cost of all processes involved (Supplementary Fig. 7), we conclude that the mechanism of D46 deprotonation studied here, involving first proton transfer from E133 to the solvent and/or galactonate and subsequent proton transfer from D46 to E133, appears feasible.

### Free energy landscapes suggest that reorientation of the empty transporter is a ratelimiting step of the cycle

Both inward- and outward-facing DgoT conformations display some flexibility; however, we did not observe transitions between these two states in single trajectories. Since a quantitative description requires multiple observations of these slow processes, we used Markov state modeling (Husic and Pande 2018) to stitch together short simulation data and analyze the transition probability between all conformational states. We featurized the trajectory data using a set of interdomain Cα atomic distances and applied time-lagged independent component analysis (tICA) to find the slowest components in the dataset. The first tICA-eigen-vector discriminates between the inward- and outward-facing conformations (Fig. 6a), and the second correlates with the (open or closed) state of the extracellular gate (Fig. 6b, Supplementary Fig. 8). Two systems were chosen as most relevant for transport: (i) galactonate-bound DgoT with protonated D46 and E133 (responsible for translocation of the substrates across the membrane) and (ii) *apo* DgoT with deprotonated D46 and E133 (responsible for reorientation of the empty transporter after substrate release). For both systems, we sampled all relevant intermediate conformations with a total of ∼26 µs (substrate-bound) and ∼35 µs (*apo*) unbiased MD simulations, and constructed Markov state models. (Fig. 6a, b). The free energy surfaces for both systems revealed a high energy barrier separating the inward- and outward-facing conformations. In the presence of galactonate, the barrier was lowered and the inward-facing state was the most energetically favorable. In contrast, the system with the *apo* protein preferred outward-facing conformations such that DgoT favors inward galactonate transport. Moreover, *apo* DgoT easily switches between outward-occluded and outward-open states, whereas substrate-bound DgoT adopts only occluded conformations. Perron-cluster cluster analysis (PCCA) (Deuflhard and Weber 2005), used to identify metastable states, revealed a difference in the direction of conformational changes between the *apo* and galactonate-bound states (Fig. 6c, d), as previously observed in the free energy surfaces (Fig. 6a, b). For substrate-bound DgoT, metastable states correspond to inward-facing, intermediate occluded, and outward-occluded states, with the highest probability for the inward conformation (Fig. 6c). For *apo* DgoT, PCCA identified inward-facing, outward-occluded, and outward-open states (lower to higher probability). The out-ward-open state was not adopted by substrate-bound DgoT: it was only adopted by *apo* DgoT, for which it represents the state with highest stationary probability (Fig. 6d). Given that substrate binding and release are fast processes compared with the conformational changes, our data show that reorientation of the empty transporter upon substrate release is the rate-limiting step in the transport cycle.

**Figure 6.**
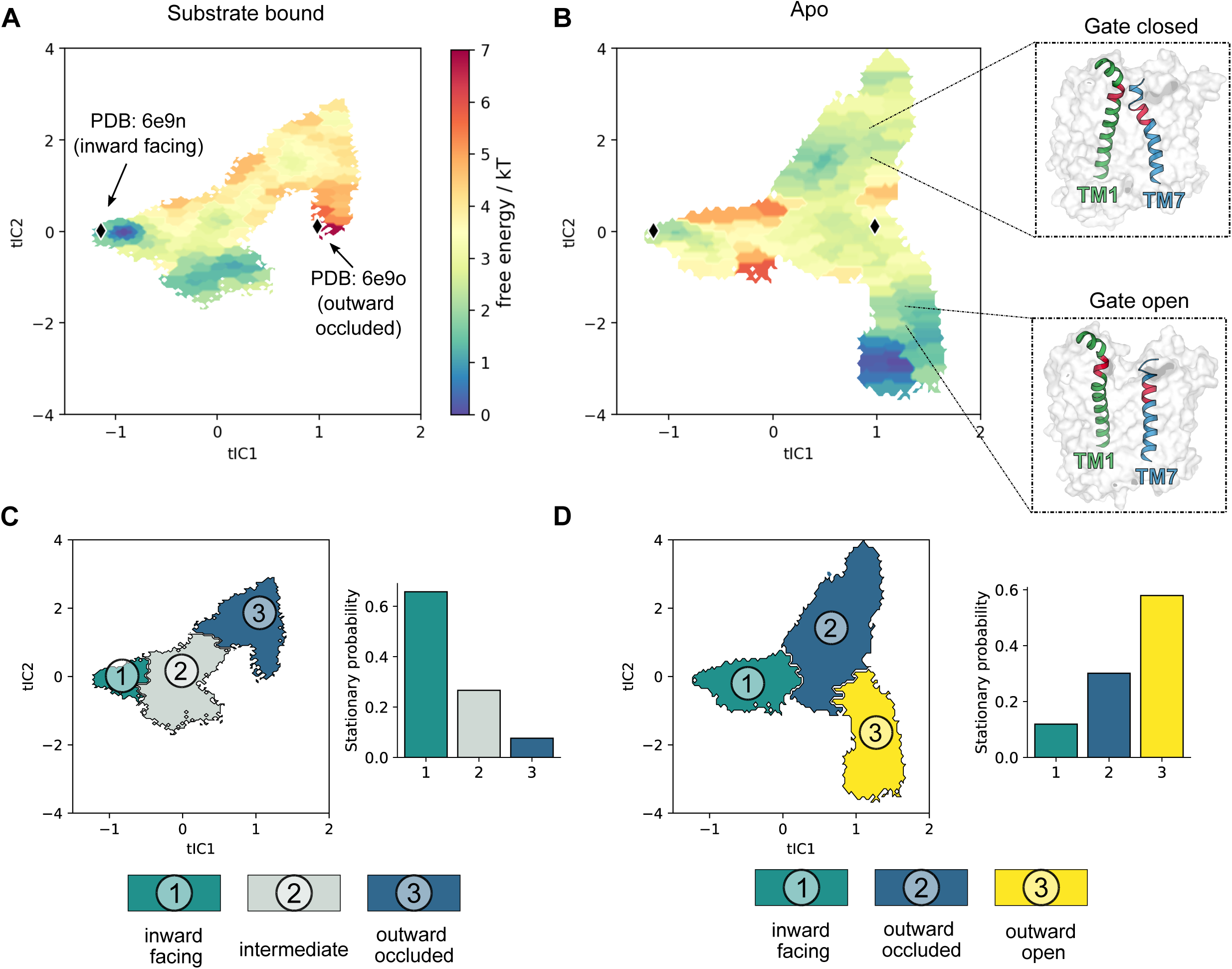
Energetical description of major conformational changes. A, B Free energy landscape for DgoT in substrate-bound (A) or *apo* (B) state. Protein conformations captured in DgoT crystal structures were projected onto the tICA space (black points). Representative snapshots illustrate how the value of tIC2 correlates with the degree of opening of the extracellular gate. C *Left:* coarse representation of intermediate metastable states obtained with PCCS for substrate-bound DgoT. *Right:* Stationaty probabilities of such metastables states. D Same representation as in (C), for apo DgoT.

### Neutralization of putative proton acceptors abolishes galactonate transport

To further investigate the function of the charged transmembrane residues, we next evaluated the effects of the corresponding neutralization mutations (D46N, E133Q, R47Q, and R126Q) on DgoT transport activity. We first monitored changes in the extracellular pH of a bacterial suspension using a micro pH electrode-based transport assay (Bosshart et al. 2019). The addition of galactonate to *E. coli* expressing WT DgoT resulted in a time-dependent increase in pH, which reached the maximum value in less than a minute. The addition of the same concentration of the galactonate epimer gluconate to DgoT-expressing *E. coli* nor of galactonate to vector-transformed (without DgoT) bacteria induced an increase in pH (Fig. 7a). All mutations tested abolished the galactonate-induced pH changes (Fig. 7b), indicating loss of transport function.

**Figure 7.**
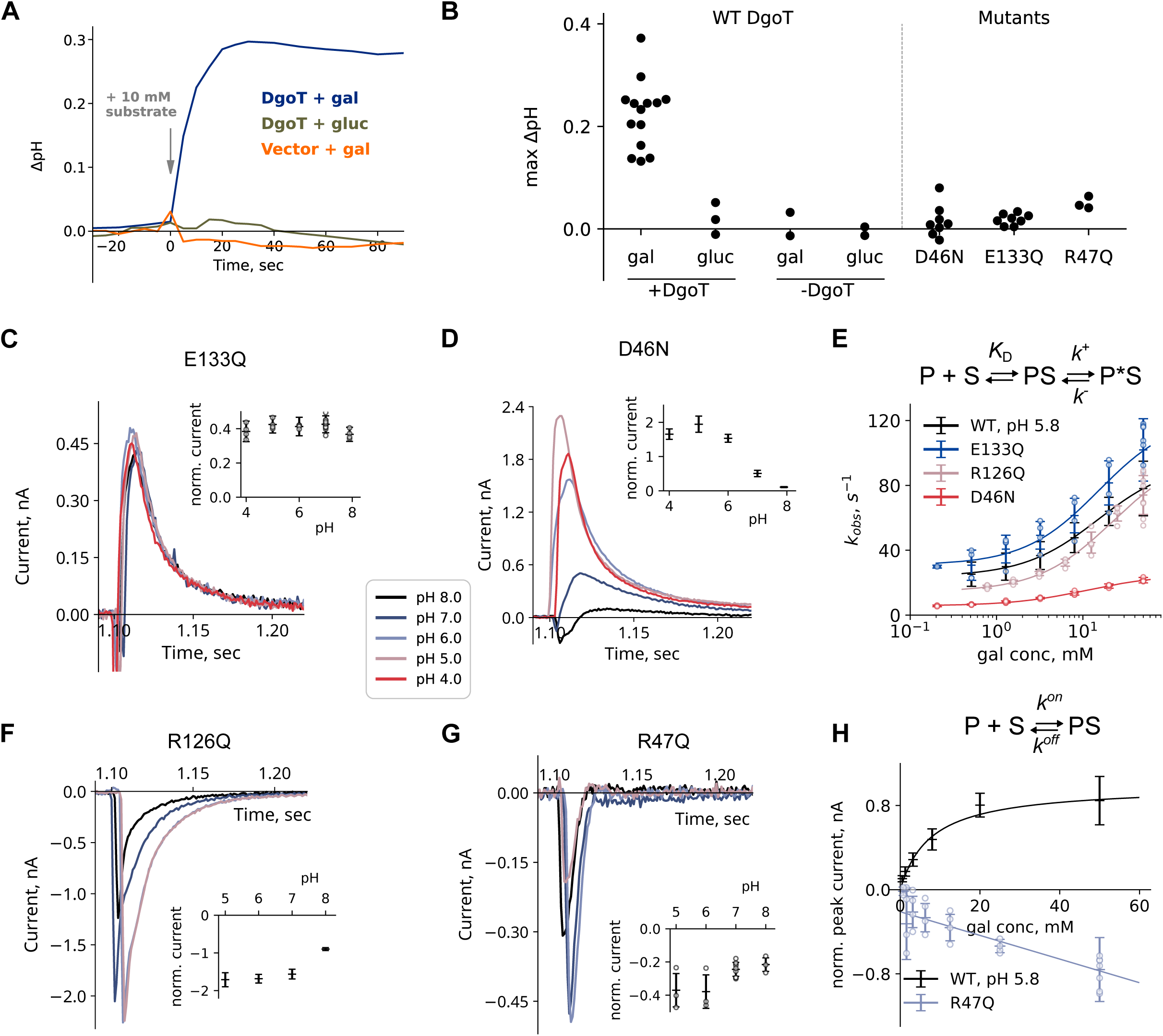
Neutralization of transmembrane charged residues abolish DgoT galactonate transport. A pH changes induced by H+-coupled galactonate, but not gluconate transport into *E.coli* cells expressing WT DgoT or transformed with the same vector without the DgoT gene. B Maximum pH changes observed in experiments with E.coli cells expressing different DgoT variants. C, D, F, G pH dependency of SSME currents initiated by galactonate concentration jump (10 mM) at symmetrical pH conditions. Experiments were performed on n=4 (C), n=5 (D), n=5 (F) and n=3 (G) independent sensors. E Concentration-dependent changes in kobs, obtained by monoexponential fit of the current decay, for WT and mutant DgoT. Solid lines are fits to a three-states induced-fit model given above the graph. Experiments were performed on n=7 (WT), n=4 (E133Q), n=4 (R126Q) and n=3 (D46N) independent sensors. H Substrate concentration dependence of the peak current values for WT and R47Q DgoT. R47Q concentration dependences were fitted to a two-state binding model given above the graph. Values of peak current values for WT DgoT (fitted to three-states model from E) are given for comparison. Experiments were performed on n=7 (WT) and n=4 (R47Q) independent sensors.

In SSME experiments, the application of galactonate elicited a positive fast pre-steady state, but not transport currents, for D46N and E133Q DgoT, indicating that these mutant transporters can bind galactonate but cannot complete the transport cycle. Galacto-nate-induced pre-steady state currents recorded with the E133Q mutant were pH independent (Fig. 7c), suggesting that protonation of this residue is responsible for the inhibition of WT DgoT at alkaline pH. At every pH tested, E133Q currents closely resembled WT currents under acidic pH conditions (Supplementary Fig. 9a), where transport is blocked by impaired proton release inside the vesicle. In contrast, D46N currents were pH dependent, with the largest peak currents observed under acidic pH conditions (Fig. 7d). Currents obtained with this mutant were biphasic (Supplementary Fig. 9b) and with slower decay than for E133Q. To determine whether the recorded currents represent the pre-steady state reaction or residual transport activity, we compared the time courses of current decay using liposomes reconstituted with different LPRs. Unlike the currents recorded with WT protein, the decay times for D46N DgoT did not systematically depend on the LPR (Supplementary Fig. 9c). Therefore, the observed reaction can be attributed to a slow pre-steady state process.

For D46N and E133Q DgoT, the peak current amplitudes changed with increasing galactonate concentrations in a saturating fashion (Supplementary Fig. 10), suggesting that these mutant transporters undergo conformational changes after rapid galactonate binding (Bazzone et al. 2021). Similar behavior was observed for WT DgoT under acidic conditions, where only the pre-steady state reaction (no transport activity) can be seen (Fig. 1a, Supplementary Fig. 3a). Fig. 7e shows that the observed rates (obtained as reciprocal decay time constants) depend on the galactonate concentration. Hyperbolic curve fitting provided the K_D_s for galactonate binding, as well as the kinetic parameters characterizing the conformational changes (Table 1). Compared with WT, conformational changes are similar for E133Q and slower for D46N but with no difference in substrate affinity. We conclude that deprotonation of D46 and E133 is necessary to complete the transport cycle; however, partial reactions that represent conformational changes can occur in the presence of galacto-nate.

**Table 1.** Overview of I_max_ values (50 mM galactonate) and kinetic parameters for transportdeficient mutants, averaged over n = 3 different sensors (errors reflect standard deviations). For R47Q analysis of current decays was limited by time resolution of the instrument, therefore only I_max_ values are given.

| Variant | $I_{\max}$ (nA) | $k_{\text{obs}}$ (s<sup>-1</sup>)<br>(50 mM galactonate) | $K_d$ (mM) | $k^-$ (s<sup>-1</sup>) | $k^+$ (s<sup>-1</sup>) |
| --- | --- | --- | --- | --- | --- |
| WT at pH 5.8 | $0.7 \pm 0.2$ | $79 \pm 14$ | 15.1 | 23.8 | 70.6 |
| D46N | $2.7 \pm 0.3$ | $68 \pm 4$ | 11.0 | 5.8 | 19.2 |
| E133Q | $1.0 \pm 0.3$ | $105 \pm 15$ | 15.3 | 30.8 | 92.2 |
| R47Q | $-0.37 \pm 0.05$ | | | | |
| R126Q | $-1.5 \pm 0.4$ | $74 \pm 11$ | 22 | 14.4 | 86.7 |

### R47 couples substrate binding and major conformational changes

For neutralizing mutations of both transmembrane arginine residues (R47Q and R126Q), galactonate application elicited fast negative currents in SSME experiments (Fig. 7f, g). For R126Q, current amplitudes were similar at pH 5–7, but lower at more alkaline pH. Peak currents changed with increasing galactonate concentrations with a hyperbolic concentration dependence (Fig. 7e), indicating conformational changes. In contrast, R47Q DgoT exhibited faster currents with amplitudes increasing linearly with increasing substrate concentration (Fig. 7h). Therefore, the observed pre-steady state reaction for R47Q differs from those observed for other transport-deficient mutants and likely represents electrogenic substrate binding rather than conformational changes (Bazzone et al. 2021).

The crystal structure of outward-facing DgoT (Leano et al. 2019), as well as our MD simulation data (Fig. 2), indicates that R47 interacts with the carboxyl group of bound galactonate. To better understand the role of this arginine residue, we carried out MD simulations with the outward-facing DgoT structure in which R47 was mutated to glutamine and D46 and E133 were protonated. In *apo* simulations, both WT and mutant DgoT assumed similar conformations with an open extracellular gate (Fig. 8a). However, galactonate failed to promote closure of the extracellular gate of R47Q DgoT (Fig. 8b). This difference is a consequence of changes in protein–substrate interactions. Galactonate is coordinated by multiple polar residues (Batarni et al. 2023) in helices from both the N- and C-domains, with the substrate located closer to TM7 in simulations with the mutant than with the WT transporter (Fig. 8c, d). Comparison of the substrate–protein interactions revealed more stable contacts for galactonate with C-domain residues such as Q264, T372, and N393 in WT; residue 47 from the N-domain interacts with the substrate only in simulations with the WT protein (not the mutant protein) (Fig. 8e).

**Figure 8.**
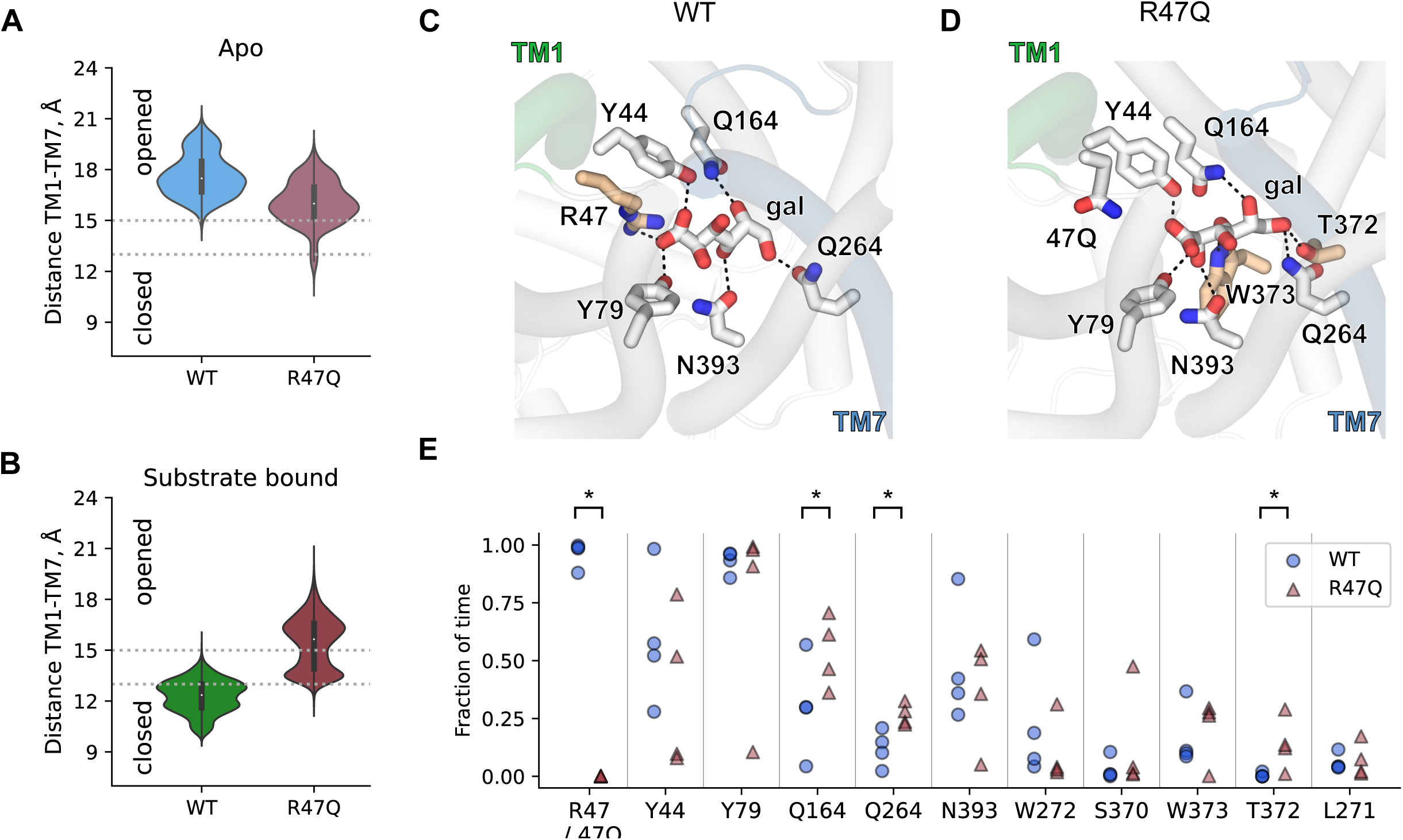
Galactonate does not induce extracellular gate closure in R47Q DgoT. A, B Probability densities for extracellular gate opening in *apo* (A) and galactonate-bound (B) simulations with WT and R47Q DgoT in outward-facing conformation with D46 and E133 protonated. C, D Position of galactonate in the binding site of WT (C) and R47Q DgoT (D). E Fraction of frames, in which distance between the side chain of specified residue and closest oxygen atom of galactonate was equal or less than 2 Å. Each data point represents one of the four independent trajectories for WT (blue points) and R47Q mutant (red points) DgoT. Significance was evaluated with the Mann–Whitney test, one-sided: \**p* < 0.05.

Our combined experimental and computational results describe substrate binding as a multiphasic process initiated by galactonate recognition and accommodation in the binding site. Subsequent direct interaction between the carboxyl group of galactonate and R47 induces closure of the extracellular gate. Both reactions are electrogenic; R47Q abolishes the second step without affecting the initial substrate binding. The fast negative component corresponding to electrogenic galactonate binding also occurs with other mutants, along with an additional slower component of positive (E133Q) or negative (R126Q) amplitude (Fig. 7c, f).

## Discussion

We combined experimental and computational approaches to describe the transport cycle of the bacterial SLC17 homolog DgoT. DgoT transport is based on an alternating access mechanism, with the transporter cycling between inward- and outward-facing states through occluded conformations in which the substrate-binding site is inaccessible from both membrane sides (Fig. 9). In the outward-facing conformation, DgoT dynamically switches between states with a closed or open extracellular gate (states 1 and 2). Protonation of both D46 and E133 stabilizes the open-gate conformation (state 2) and permits galactonate binding from the periplasmic side (state 3), followed by closure of the extracellular gate. Formation of the outward-occluded conformation is the initial step of major conformational change (state 4) that brings the protein into an inward-facing conformation (states 5, 6, and 7). In this conformation, deprotonation of D46 (state 5) results in an open intracellular gate that permits galactonate release (state 6). D46 deprotonation may occur via multiple pathways, as shown for proton transfer in other transporters (Swanson 2022). Our multiscale simulations demonstrate that D46 deprotonation is possible via initial proton release from E133, either to the intracellular solvent or to galactonate, followed by H^+^ transfer from D46 to E133 (state 6). Galactonate can unbind in a protonated or non-protonated form; protonation might promote galactonate release by weakening the electrostatic interaction with R47 and stabilizing intracellular gate opening. H^+^ transfer from D46 to E133 (state 6) and subsequent release from E133 results in intracellular gate closure (state 7); reorientation of the empty transporter to outward-facing conformation completes the cycle (state 8).

**Figure 9.**
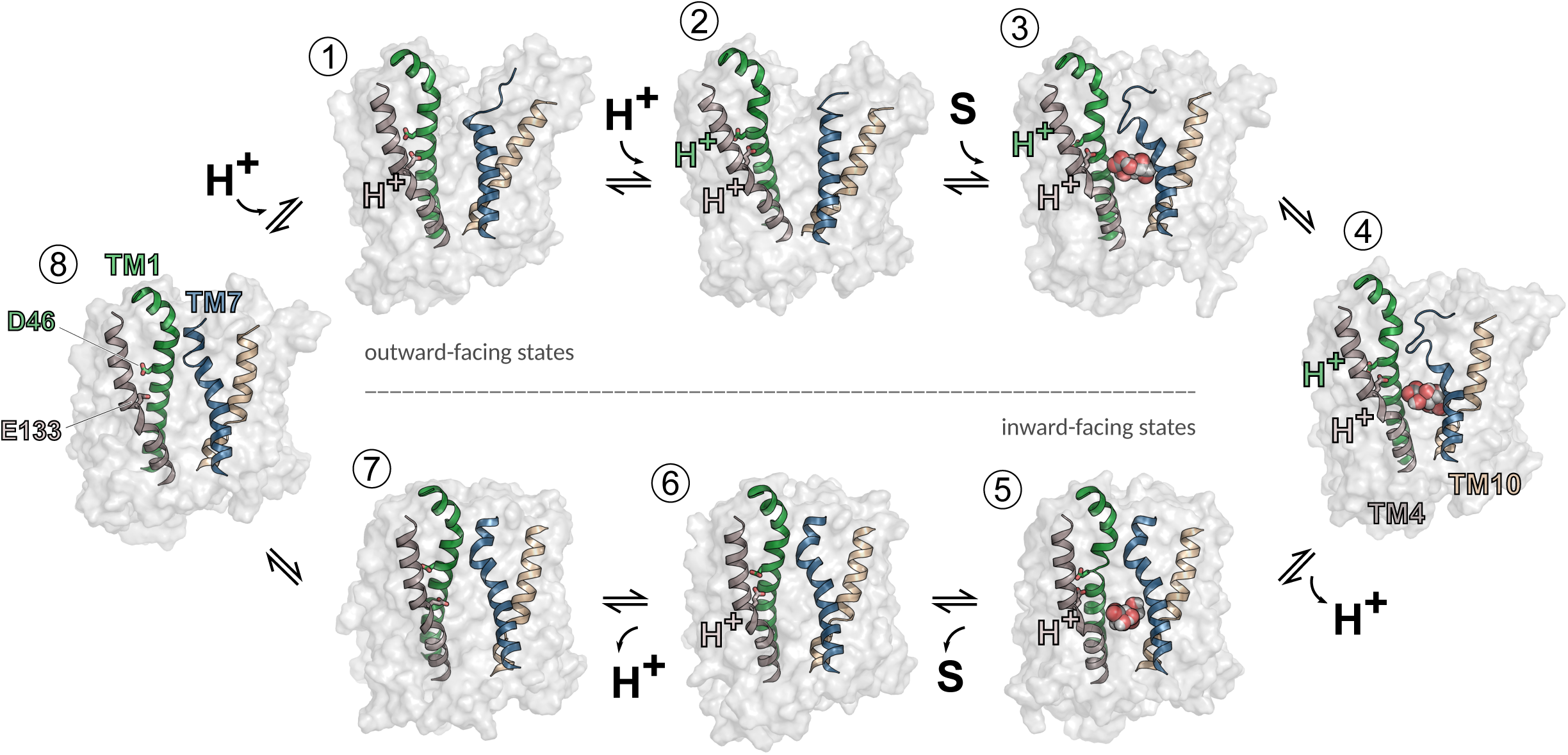
Schematic transport cycle of DgoT. In outward-facing conformation (1) extracellular gate opening is stabilized by protonation of D46 and E133 (2). Galactonate binding (3) induces major conformational changes (4). In the inward-facing state (5) proton release from D46 leads to galactonate release to the intracellular solution (6), followed by the release of the second proton (7) and reorientation of the empty carrier to outward-facing conformation (8).

Our classical MD simulations revealed how the protonation state of the two acidic transmembrane residues is coupled to the conformational changes necessary for substrate binding and release. In the *apo* state, deprotonated D46 and deprotonated E133 both serve as “latches” by interacting with R126 (TM4) and R47 (TM1), respectively; this strategy is commonly used by MFS transporters (Drew et al. 2021). Protonation of the two acidic residues unclasps these latches, albeit with distinct effects on extracellular gate mobility (Fig. 2d and Fig. 3d). For D46, H^+^ association releases R126 and shifts the equilibrium toward a state with an open extracellular gate, whereas protonation of E133 alone stabilizes the gate in an only slightly opened state. Double protonation is necessary for full opening of the extracellular gate, which makes substrate binding most effective (Fig. 2d and Fig. 3d). Galactonate binding promotes closure of the extracellular gate for all four protonation state combinations (Fig. 2e). However, stable gate closure requires protonation of E133 (Fig. 2e), which disrupts the E133–R47 salt bridge and gives TM1 the flexibility to come into closer contact with TM7 upon galactonate binding (Fig. 2c).

Arginine residues at positions 47 and 126 are conserved within SLC17 transporters (Supplementary Fig. 2). R47 forms an electrostatic interaction with galactonate in the binding site that is suggested to be responsible for its recognition (Leano et al. 2019). However, R47Q DgoT exhibits galactonate-specific pre-steady state currents (Fig. 7g) that represent a binding reaction rather than rate-limiting conformational changes. Therefore, our results assign a role beyond substrate recognition to R47: this arginine connects the substrate-binding site to the protonation site and triggers closure of the extracellular gate, much like H322 in lactose permease LacY (Kumar et al. 2015). Pre-steady-state currents of R126Q DgoT indicate electrogenic conformational changes rather than a binding reaction. R126 might be involved in tuning the pKa of D46 at different stages of the transport cycle or be necessary for conformational changes in *apo* DgoT.

The proposed transport mechanism (Fig. 9) fully agrees with all published experimental data. It accounts for the stoichiometrically coupled symport of two H^+^ and one galactonate, as determined experimentally (Fig. 1e). Solubilized DgoT does not bind protons after the addition of galactonate (Fig. 1f), as would be expected if protonation of the transporter precedes substrate binding. Neutralization of residue D46 or E133 prevents coupled galac-tonate transport, as expected for the removal of obligatory protonation sites. Our transport scheme can also account for recently reported experiments demonstrating that E133Q, but not D46N, DgoT is capable of galactonate exchange (Batarni et al. 2023). The E133Q mutation has no effect on galactonate release and substrate exchange: it exclusively abolishes forward transport by preventing *apo* translocation back to the outward-facing conformation. In contrast, D46N prevents forward transport as well as galactonate exchange, since depro-tonation of D46 is required for galactonate release (Fig. 4). The surprising result that the additional E133Q mutation restores galactonate exchange in D46N/E133Q DgoT might be explained if galactonate is predominantly released in a protonated form by WT DgoT. In D46N DgoT, galactonate may still be protonated via H^+^ transfer from E133 although the closed intracellular gate would prevent its release (Fig. 4d). In D46N/E133Q DgoT, galact nate cannot be protonated and the slightly higher open probabilities for the intracellular gate with non-protonated galactonate make the double mutant competent for exchange.

1. *E. coli* can use galactonate as the sole carbon source (Deacon and Cooper 1977), demonstrating that DgoT transports as effectively as alternative glucose transporters, such as lactose permease LacY (Kaback and Guan 2019) or xylose symporter XylE (Madej et al. 2014). LacY and XylE transport uncharged sugar molecules with a stoichiometry of 1 proton:1 substrate, whereas DgoT co-transports two protons with the negatively charged galac-tonate, together resulting in a positive net transported charge. Most likely, the adjusted transport stoichiometry permits DgoT to utilize ΔpH and ΔΨ and provides high driving forces, thus optimizing bacteria for nutrient uptake when resources are limited.

Transport substrates and stoichiometries vary substantially among MFS transporters. LacY (Kaback and Guan 2019) and fucose transporter FucP (Dang et al. 2010) share a stoichiometry of 1 H^+^:1 substrate. In LacY, protonation of a single site, E325 (equivalent to E133 in DgoT (Kaback and Guan 2019)), permits lactose binding and transition to the occluded conformation. FucP has two protonation sites, D46 and E135, that are homologous to D46 and E133 in DgoT. However, only D46 is accessible from the extracellular solution, with E135 serving only as part of the proton transfer pathway. SLC17A5/sialin differs from DgoT in having broad substrate specificity: it recognizes various sialic acids via electrostatic interaction with two conserved arginines, R168 and R57. Two glutamates, E171 and E175, serve as protonation sites. Protonation of E171 releases R168 and permits its interaction with the substrate (Hu et al. 2023). Subsequent H^+^ transfer to E175 is followed by substrate transfer to a more cytoplasmic binding site close to R57 and translocation to the inward-facing conformation. Such coupling of substrate translocation to proton transfer results in the electroneutral co-transport of one H^+^ and one sialic acid molecule.

The comparison of DgoT with FucP and SLC17A5/sialin reveals how small variations in the arrangement of protonation sites can adjust transport stoichiometries of H^+^-coupled transporters with conserved architecture and transport mechanisms. The SLC17 family en-compasses organic anion uniporters (Ishikawa et al. 2008; Iharada et al. 2010), H^+^-gluta-mate exchangers (Kolen et al. 2023), and a H^+^-sialic acid symporter (Hu et al. 2023). Certain SLC17 transporters can also function as a Na^+^-PO_4_^3-^ symporter (Iharada et al. 2010; Preobraschenski et al. 2018). Therefore, our work may serve as a framework to understand the mechanisms underlying the diversity of SLC17 transport mechanisms.

## Methods

### Expression and protein purification

The full-length DgoT gene (GenBank accession number AKK15832.2) was subcloned into a pQE60 vector through the NcoI and HindIII restriction sites in fusion with a C-terminal thrombin cleavage site and decahistidine tag. Mutant constructs were generated using PCR-based mutagenesis and verified by DNA sequencing. Protein expression and purification were performed using a published procedure, with modifications (Leano et al. 2019). *E. coli* C41 cells transformed with pQE60 DgoT WT were grown at 37 °C in TB medium supple- mented with 2 mM MgSO_4_. When an OD_600_ of 0.6–0.8 was reached, gene expression was induced with 1 mM IPTG and cells were grown for a further 4 h (typical yield of 15 g per 1 L culture). After sedimentation, cells were flash frozen in liquid nitrogen and stored at -80 °C for later use, Next, cells (15 g) were resuspended in 20 mM Tris (pH 7.4) and 300 mM NaCl (50 mL volume) containing complete protease inhibitor cocktail (Roche) and lysed by soni- cation. Debris was removed by centrifugation at 12,000 × *g* for 15 min, and membranes were collected at 200,000 × *g* for 1 h, flash frozen in liquid nitrogen, and stored at -80 °C until use.

A frozen membrane pellet from 15 g cells was resuspended in 20 mL membrane buffer (20 mM Tris (pH 7.4), 150 mM NaCl) containing cOmplete protease inhibitor cocktail, using a glass Dounce homogenizer, and then *n*-dodecyl-D-maltoside (DDM) was added to 1.4%. Membranes were solubilized for 2 h at 4 °C and the insoluble fraction was removed by ultracentrifugation at 75,000 g for 30 min at 4 °C. The supernatant was diluted 1:2 with solubilization buffer, imidazole was added to 15 mM, and pH was adjusted to 7.8–8.0. CoNTA resin (3 mL) was washed with 10 column volumes (CV) of membrane buffer, added to the supernatant, and incubated at 4 °C for 1.5 h under gentle agitation. The resin was then washed with 10 CV of wash1 buffer (20 mM Tris, 150 mM NaCl, 15 mM imidazole, 0.1% DDM, pH 8.0) and 10 CV of wash2 buffer (20 mM Tris, 150 mM NaCl, 0.1% lauryl maltose neopentyl glycol (LMNG), pH 7.4). Protein was eluted with 4 CV of elution buffer (20 mM Tris, 150 mM NaCl, 0.1% LMNG, 150 mM imidazole, pH 7.4) into 0.5 CV fractions, and 5 mM EDTA was added immediately after collection. Protein concentration in the eluted fractions was estimated by measuring the absorbance at 280 nm (NanoDrop), and fractions with the highest protein concentrations were combined. Imidazole was removed using a PD-10 desalting column, with protein eluted with buffer (20 mM Tris, 150 mM NaCl, 1 mM EDTA, 0.05% LMNG, pH 7.4) and stored overnight at 4 °C.

The purified protein was concentrated to 5–8 mg/ml using a 50 kDa molecular weight cutoff (MWCO) centrifuge concentrator (Millipore) and loaded in 0.5 mL portions onto a Superdex 200 Increase 10/300 GL column (GE Healthcare Life Sciences) preequilibrated with size-exclusion chromatography (SEC) buffer (20 mM HEPES, 150 mM NaCl, 0.05% LMNG, pH 7.4). The peak fractions were combined, flash frozen in liquid nitrogen, and stored at - 80 °C until use.

### Reconstitution of proteoliposomes

1. *E. coli* polar lipids (Avanti Polar Lipids, *E.coli* polar lipid extract, 25 mg/ml solution in chloroform) were dried under nitrogen and then under vacuum overnight. The dried lipid film was resuspended to a lipid concentration of 10 mg/ml by stirring in reconstitution buffer (1 mM HEPES pH 7.4, 150 mM NaCl, 2 mM MgSO_4_) for 1 h at room temperature (RT). After the lipids were completely dissolved, the suspension was frozen in liquid nitrogen, stirred for another 1 h at RT, and sonicated using a UP50H ultrasonic processor equipped with a microtip until clear (2 or 3 cycles of 30 s). The formed liposomes were destabilized by adding 0.6% Triton X-100 and incubated for 45 min at RT under gentle agitation. Purified DgoT was added to the destabilized liposomes at a LPR of 5, 10, or 20 and then incubated for 1 h at 4 °C. To remove the detergent, 150 mg SM2 Bio-beads was added per 1 mL liposome suspension. After incubation for 1 h at 4 °C, another 150 mg Bio-beads was added per 1 mL suspension, followed by another 1 h incubation at 4 °C, and then the beads were removed using a disposable column. A third volume of Bio-beads was then added (400 mg per 1 mL suspension), incubated overnight at 4 °C, and removed using a disposable column. Empty liposomes were prepared in parallel using the same procedure but with no added protein.

### pH electrode-based transport measurements

DgoT was expressed in *E. coli* C41 cells as described for protein purification. After protein expression, cells were pelleted, washed twice in assay solution (250 mM KCl, 1 mM MgSO_4_, 2 mM CaCl_2_), and resuspended in assay solution to an OD_600_ 15. All centrifugations were performed at 2,200 × *g* for 7–8 min, and cells were resuspended by vortexing. Substrates to be assayed were dissolved in assay solution at a concentration of 160 mM and the pH was adjusted to 6.5–6.7 with KOH such that the substrate pH was lower than the pH of the bacterial suspension for each experiment. A 800 μL volume of bacterial suspension was transferred to a 2 ml reaction tube (Eppendorf) and the pH of the suspension (i.e., of the extracellular medium) was measured using a micro pH electrode with an integrated temperature sensor (Xylem, SI Analytics) under constant stirring and adjusted to pH 6.7 using KOH and HCl. After recording the baseline for 60 seconds, 50 µL of the corresponding compound was added to the bacterial suspension (final concentration: 10 mM). All experiments were performed at 20–21 °C. For experiments with DgoT in detergent, purified DgoT (stored at - 80 °C) was thawed and SEC buffer was replaced using a PD-10 desalting column with assay solution containing 150 mM NaCl and 0.05% LMNG (pH ∼7.4). Purified protein was used at a concentration of 4 µM in all experiments. The experiment was repeated six times, with comparable results.

### SSM-based electrophysiology

Gold electrode sensors (1 or 3 mm) were prepared as previously described (Bazzone et al. 2017). Briefly, sensors were incubated for at least 30 min in an octadecane thiol solution, and then rinsed thoroughly with isopropanol and water. The solid-supported membrane (SSM) was prepared by pipetting 1.5 µL diphytanoyl phosphatidylcholine dissolved in *n*-decane onto the electrode surface, followed by 100 µL aqueous buffer. Immediately prior to measurements, liposome samples were thawed, diluted to a final lipid concentration of 1 mg/mL and briefly sonicated. Each liposome sample (10 µL) was pipetted onto a SSM sensor and adsorbed by centrifugation at 2,200 × *g* for 30 min at RT. All experiments were repeated for at least three sensors, with each condition measured at least twice. All solutions were buffered in 100 mM potassium phosphate (KPi) for each pH used.

For measurements with a single solution exchange protocol, three phases of 1 s duration were applied: flow of nonactivating (NA) solution, activating (A) solution, and NA solution. Only the A solution contained galactonate. In experiments with variable pH, pH of NA and A solutions was kept constant within a single experiment. Between experiments using different pH values, the sensor was incubated at the new pH for 5 min to equilibrate the intraliposomal pH. In experiments with WT, D46N, E133Q, and R126Q DgoT, the NA solution contained gluconate to compensate for galactonate in the A solution. For R47Q DgoT, glutamate was used in NA solution instead, allowing a direct comparison of the responses to galactonate and gluconate applications.

To determine the apparent pK values, normalized peak currents (measured using 3 mm sensors) were fitted with one of the following equations:

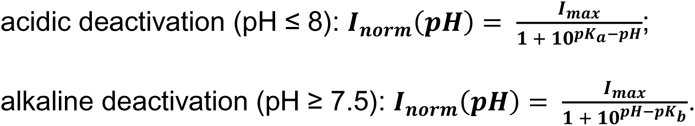

Currents were normalized to the *I_max_* value obtained by the fit of peak currents measured on the same sensor.

For the analysis of pre-steady state currents, 1 mm sensors were used. Rate constants for the observed charge displacements (k_obs_= 1/τ) were derived from the transient currents by fitting the decay with a monoexponential function I = A * exp(-t/τ). This two-step reversible reaction can be described as follows:

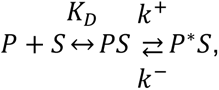

where *P* is the protein, *S* is the substrate, and *P\** is the protein after conformational changes. The first step is the binding reaction described by the dissociation constant K_D_, and the second step is the substrate-induced conformational change characterized by the forward and reverse rate constants. Assuming that substrate binding is rapid, the observed rate constant has hyperbolic dependence on the substrate concentration (Smirnova et al. 2006):

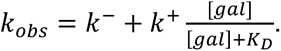

Measurements under asymmetrical pH conditions were done using a double solution exchange configuration. A resting (R) solution phase of 1 s and incubation period of 5– 20 min were added to the beginning of each measurement to allow the intraliposomal pH to adjust to the pH of the R solution. Afterwards, a normal single solution exchange protocol was used to establish the pH gradient (during NA phase) and substrate gradient (A phase). To determine the transport stoichiometry, we used a reversal assay as previously described (Thomas et al. 2021), with a single solution exchange protocol comprising three phases (NA, A, and NA) extended to 2, 3, and 3 s, respectively. Between measurements, the sensor was rinsed 3–5 times with NA solution; the current responses were recorded and used as a baseline. The same protocol was used for samples with empty liposomes to account for solution exchange artifacts. The entire A phase was integrated to obtain the transported charge values. After subtraction of the negative control (integrated current recorded with empty liposomes), the transported charge values were used to determine the transport stoichiometry. The internal (NA) solution used for this experiment had a pH of 7.3 and contained 0.5 mM galactonate and 7.5 mM gluconate, and applied external (A) solutions had a pH of 7.6 and contained X mM galactonate and 8–X mM gluconate. This setup resulted in dµ _gal_/ dµ _H+_ values of 0–4.

### MD simulations

DgoT protein coordinates obtained from the Protein Data Bank (inward-facing, PDB ID: 6E9N; outward-facing galactonate-bound, PDB ID: 6E9O (Leano et al. 2019)) were used as the starting coordinates for MD simulations. The D46 and E133 protonation state and substrate occupation were modified as described in the text. Standard protonation states at neutral pH were assigned to all other residues (deprotonated aspartate and glutamate residues, singly protonated histidine residues) except for H56, which forms a salt bridge with E180 and, therefore, was set as doubly protonated. Missing residues (235–242 in the in-ward-facing structure, 231–243 and 277–290 in the outward-facing structure) were modeled using the SWISS MODEL server (Waterhouse et al. 2018) with a final residue range of 27– 442 used for both protein structures. The N- and C-termini were capped with neutral acetyl and methylamide groups, respectively. For modeling DgoT single point mutations, we used PyMOL (Schrödinger, LLC 2015).

The initial protein orientation within the membrane was set to the corresponding DgoT structure available in the Orientations of Proteins in Membranes database (Lomize et al. 2012). The protein was then embedded into a phosphatidylcholine (POPC) bilayer using *g_membed* (Wolf et al. 2010) in GROMACS and solvated in a box with dimensions ∼120 × 120 × 100 Å, which was chosen to ensure a minimum distance between periodic copies of at least 30 Å. The simulation temperature was 310.15 K. The protein/membrane system was surrounded by a ∼100 mM solution of Na^+^ and Cl^−^ ions. Ions were described using default CHARMM parameters, and the CHARMM TIP3P model was used for water molecules. MD simulations were performed using the GROMACS software package (versions 2018, 2020, and 2022) (Abraham et al. 2015) with a CHARMM36m force field (Huang et al. 2017). Galactonate parameters were obtained using the SwissParam server (Zoete et al. 2011) and added to the forcefield. An integration time step of 2 fs was used. Van der Waals interactions were calculated with the Lennard–Jones potential and a cutoff radius of 1.2 nm, with forces smoothly switched to zero in the range of 1.0–1.2 nm and no dispersion correction applied. Electrostatic interactions were calculated by the particle mesh Ewald method (Essmann et al. 1995), with a real-space cutoff distance of 1.2 nm. All simulations were done in an isothermal–isobaric ensemble, with the temperature set to 310 K using a v-rescale thermostat (Bussi et al. 2007) and a time constant of 0.5 ps. The thermostat was applied separately to the protein, lipid bilayer, and aqueous solution containing ions. The same groups were used for the removal of the center-of-mass linear motion.

The protein was equilibrated in three steps using the velocity-rescale thermostat and Berendsen (Berendsen et al. 1984) pressure coupling. The first step lasted 50 ns and was run with positional restraints on protein atoms with a harmonic potential with a force constant of 1000 kJ mol^−1^ nm^−2^ to allow for equilibration of water and ions. In the second step, only the backbone atoms of the protein were restrained to enable side chains to equilibrate. Lastly, the system was equilibrated for 1 ns without positional restraints to obtain the velocities used in the following production runs. Production MD simulations used Parrinello–Rahman (Parrinello and Rahman 1981) pressure coupling in a semi-isotropic manner with a time constant of 0.5 ps.

### Classical WT-MTD simulations

The PLUMED v2.8.1 plugin (Tribello et al. 2014) interfaced with GROMACS v2020.4 (Abraham et al. 2015) was used for simulations, along with MD settings similar to those described for MD simulations. The collective variable used to describe the conformational change of D46 is the N-Cα-Cβ*-C*γ dihedral angle (χ_1_). WT-MTD parameters were used as previously reported (Chiariello et al. 2020; Chiariello et al. 2021): Gaussian height, 1.2 kJ/mol; Gaussian sigma, 0.35 rad; bias factor, 15; and Gaussian deposition frequency, 500. Employing a previously described algorithm (Tiwary and Parrinello 2015), we applied reweighting to the dihedral angle serving as the CV to reconstruct the histogram of the sampled configurations. WT-MTD simulations were run for 100 ns, which was sufficient to achieve a converged free energy surface (shown in Supplementary Fig. 11).

### QM/MM simulations

QM/MM simulations were carried out using GROMACS v2020.4 (Abraham et al. 2015) and CPMD v4.3 (Hutter et al. 2000), coupled with MiMiC v0.2.0 (Olsen et al. 2019; Bolnykh et al. 2019). MiMiC-QM/MM input files were generated using MiMiCPy v0.2.1 (Raghavan et al. 2023). Different QM partitions were used to investigate the different processes. Region QM1 was used to simulate proton transfer from protonated E133 to the immediate neighbor water molecule and, therefore, encompassed the sidechains of E133, the galactonate molecule, and the three water molecules bridging these. The initial QM/MM configuration was obtained from WT-MTD simulations of the system in which both D46 and E133 were protonated. A snapshot in which E133 and galactonate were connected through water molecules was identified by visual inspection. Region QM2 included the hydronium ion formed by proton release from E133, the substrate galactonate, and the first layer of surrounding water molecules, enabling the simulation of proton migration to either galactonate or solvent. The initial QM/MM configuration was taken from the window corresponding to the products (CV1 = 0.35 Å) of the first TI proton transfer simulation (see section “TI simulations of proton transfer”). Lastly, QM3 included the sidechains of E133 and D46 to enable the study of proton transfer from protonated D46 to deprotonated E133. Here, the initial QM/MM configuration was extracted through visual inspection from the WT-MTD simulation of the system with protonated D46 and now-deprotonated E133 in order to identify a snapshot with D46 in the *close* conformation and H-bonded to E133. The total number of QM atoms for each partition was 44 (QM1), 82 (QM2), and 18 (QM3).

The QM problem was solved using the DFT (Hohenberg and Kohn 1964; Kohn and Sham 1965) using the BLYP functional (Becke 1988). Core electrons were described using norm-conserving pseudopotentials of the Martins−Troullier type (Troullier and Martins 1991), and the valence electrons were treated explicitly. Monovalent carbon pseudopotentials were used to saturate the dangling bonds at the boundaries between the QM and MM regions (von Lilienfeld et al. 2005). Isolated system conditions were achieved using the method of Martyna and Tuckerman (Martyna and Tuckerman 1999). The MM system was treated with the CHARMM36m force field (Klauda et al. 2010) for protein, lipid, and ions, and the TIP3P model for waters, as in the classical MD simulations. Electrostatic interactions between the QM and MM subsystems were described using the Hamiltonian electrostatic coupling scheme (Laio et al. 2002; Olsen et al. 2019). The short-range electrostatic interactions between the QM partition and any MM atom within a cutoff of 30 a.u. from the QM region were explicitly considered, with a seventh-order multipole expansion of the QM electrostatic potential used for long-range interactions.

Constant temperature simulations were carried out at 300 K using a Nosé–Hoover thermostat (Nosé 1984; Hoover 1985) (with a coupling frequency of 3500 cm^-1^).

Initially, all systems underwent relaxation through simulated annealing, with gradual reduction in the temperature of the QM/MM system by removing excess kinetic energy in each step (velocity multiplied by a factor of 0.99). Subsequently, systems were linearly reheated to 300 K and the target temperature increased using a Berendsen thermostat coupled to the system (coupling strength of 5000 a.u.). A Born–Oppenheimer molecular dynamics (BOMD) scheme was used. A timestep of 10 a.u. (∼ 0.24 fs) for the initial geometry optimization using an annealing protocol. Finally, systems with QM1 and QM3 partitions underwent 20 ps of QM/MM MD equilibration at 300 K with a timestep of 20 a.u. (∼0.48 fs) before running TI simulations; details of the QM/MM MD simulations with the QM2 partition are provided in the section “QM/MM MD proton migration simulations”.

### TI simulations of proton transfer

The free energy profiles of the two proton transfers investigated here was determined through TI in its Blue Moon sampling implementation (Ciccotti and Ferrario 2004; Ciccotti et al. 2005). The collective variable (CV) used to describe proton transfer from E133[H] to its adjacent water molecule (CV1) was the difference between (i) the distance between the carboxylate oxygen atom and the transferred proton of E133[H] and (ii) the distance between the same proton and the oxygen atom of water. Similarly, the CV2 used to simulate proton transfer from D46[H] to E133 was the difference between (i) the distance between the carboxylate oxygen atom and the transferred proton of D46[H] and (ii) the distance between the same proton and the carboxylate oxygen atom of E133.

A total of 17 and 15 independent constrained MD simulations were conducted for each TI calculation, respectively, with windows separated by 0.08 Å increments. At each CV value (either CV1 or CV2), the systems were simulated for 2 ps and the last snapshot was taken to set up the next window. The constraint force at each CV value was calculated by averaging the Lagrange multiplier of the Shake algorithm after discarding the first 0.5 ps (i.e., when it reached convergence). The free energy profiles were then obtained by integrating the constraint force along the CV using the trapezoid method. For both TI profiles, the correction term was found to negligible because fluctuations in the angle connecting donor, proton, and acceptor are minimal.

### QM/MM MD proton migration simulations

Starting with the final structure (CV1 = 0.35 Å) obtained from the first TI proton transfer simulation, in which the proton was transferred from E133[H] to the adjacent water molecule, we ran a 40 ps QM/MM MD simulation with a timestep of 20 a.u. (∼0.48 fs), maintaining the same constraint applied during TI. This served to sample an initial set of structures of the solvated hydronium ion forming an ion pair with negatively charged E133. We then applied a cluster analysis approach utilizing the *cluster* module of GROMACS. E133, D46, galactonate, and the first layer of surrounding water molecules (within 4 Å) were considered in the RMSD calculation, and clustering was performed with the *gromos* method(Daura et al. 1999) and a 0.04 Å cutoff. This approach resulted in the identification of nine distinct clusters. Subsequently, seven representative snapshots, corresponding to the centroids of the seven most populated clusters and covering over 95% of the QM/MM MD trajectory, were selected as starting configurations for the QM/MM MD simulations. Notably, these structures are characterized by variations in their hydrogen bond networks and the number of connecting water molecules among E133, hydronium ion, and galactonate. Statistical information pertaining to the proton transfer dynamics of these QM/MM MD simulations is presented in Supplementary Table 2.

### Quantification of electrostatics

Electrostatics was quantified with g_elpot (Kostritskii et al. 2021; Kostritskii and Machtens 2023); source code, installation instructions, and usage recommendations can be found at https://jugit.fz-juelich.de/computational-neurophysiology/g_elpot. The distribution of electro-static potential was calculated via the smooth particle mesh Ewald (SPME) method. For our system, the SPME potential was calculated on a grid of 256 × 256 × 208 points with an inverse Gaussian width β of 20 nm^-1^. We exploited the extended functionality of the tool (Kostritskii and Machtens 2023) to calculate the potential for the carboxyl groups of protonated D46 and E133. The electrostatic potential is an average of the SPME potential in a 0.15-nm sphere around the following atom groups (CHARMM naming convention): OD1, OD2, and CG of D46 and OE1, OE2, and CD of E133. Since two selected groups are chemically identical, the residue-specific short-range part of the potential is the same and, therefore, comparison of the potential values is justified. The time course of the electrostatic potential was averaged across each trajectory.

### Markov state model construction

Markov state models (MSMs) were constructed from multiple unbiased MD trajectories using PyEMMA 2 software (Scherer et al. 2015). To better capture the movement of N- and C-terminal domains relative to each other, we selected every 5th Cα atom of the TM helices and measured the distances between each selected atom located on the N-terminal (TM1– 6) and C-terminal (TM7–12) domains, respectively. This resulted in 27 × 26 = 2809 pairwise distances. To reduce the dimensionality, we used a time-lagged independent component analysis (tICA) with a lag time of 50 ns on a set of unbiased MD simulations to obtain the slowest collective motions. For a description of the major conformational changes, we kept the first two independent components (ICs) because they had slower timescales than the other ICs (Supplementary Fig. 11b, c). Since the starting point of the unbiased simulations was the corresponding crystal structure (either inward- or outward-facing), only regions of constructed conformational space that are close to the starting conformations were sampled. To overcome the sampling gap between unbiased simulations that started with different crystal structures, we identified conformations that were closest to the central region of conformational space and, therefore, should represent different occluded states and initiated new unbiased simulations from them. This process was repeated iteratively until the implied timescales converged. A total of ∼25 µs (double protonated substrate-bound DgoT) and ∼37 µs (DgoT *apo*) simulation data were obtained and used for further data analysis, with individual trajectories of 400–800 ns in length (Supplementary Fig. 11a). The implied time- scales plot demonstrates the Markovian behavior after a lag time of ∼ 80 ns (Supplementary Fig. 11b, c). For constructing the MSM, a lag time of 100 ns was chosen from the implied timescales plot. For model validation, we performed a Chapman–Kolmogorov test with three metastable states for both substrate-bound and *apo* DgoT systems (Supplementary Fig. 11d, e). Metastable states were identified using Perron-cluster cluster analysis (PCCA) (Deuflhard and Weber 2005). For each system, the implied timescale plots (Supplementary Fig. 11b, c) revealed two slow processes; therefore, we clustered the microstates into three metastable states.

### Data analysis and statistics

Each experimental dataset represented as means ± error bars was generated by performing the same experiment on multiple sensors (sensor numbers are given in the respective legends). Error bars represent standard deviations of the average value, and individual data points are given for n < 5. Hydration profiles (Fig. 3c and 4c) were calculated as the number of water molecules in 2 Å sections along the z-coordinate in each frame; average values and standard deviations are plotted. Violin plots were used to visualize the shape of data distribution and highlight trends. For statistical analysis of two groups (Fig. 8e), we used Mann–Whitney test, one-sided; significance is indicated as *p < 0.05.

### Data Availability

This study includes no data deposited in external repositories

## Acknowledgement

We thank Drs Andre Bazzone, Bassam Haddad, Andrei Kostritskii, Piersilvio Longo and Jan-Philipp Machtens for helpful discussions and Meike Berndt for excellent technical support. This work was supported by the Deutsche Forschungsgemeinschaft (German Research Foundation) to Ch.F. (FA 301/15–2), P.C. (CA 973/27-2) and M.A.P. (AL 2511/1-2) as part of Research Unit FOR 2518, DynIon. The authors gratefully acknowledge computing time on the supercomputer JURECA at Forschungszentrum Jülich under grants dgoth and vglut-pt.

## Author contributions

N.D., P.C., M. A.-P., and Ch.F. conceived the study. P.C., M. A.-P., and Ch.F. supervised the project, D.N. designed, performed and analyzed experiments, D.N., S.G. and C.A. designed, performed and analyzed simulations, N.D. and Ch.F. wrote the manuscript with inputs from other authors.

