## Supplementary Fig. 1 for "Transport mechanism of DgoT, a bacterial homolog of SLC17 organic anion transporters"

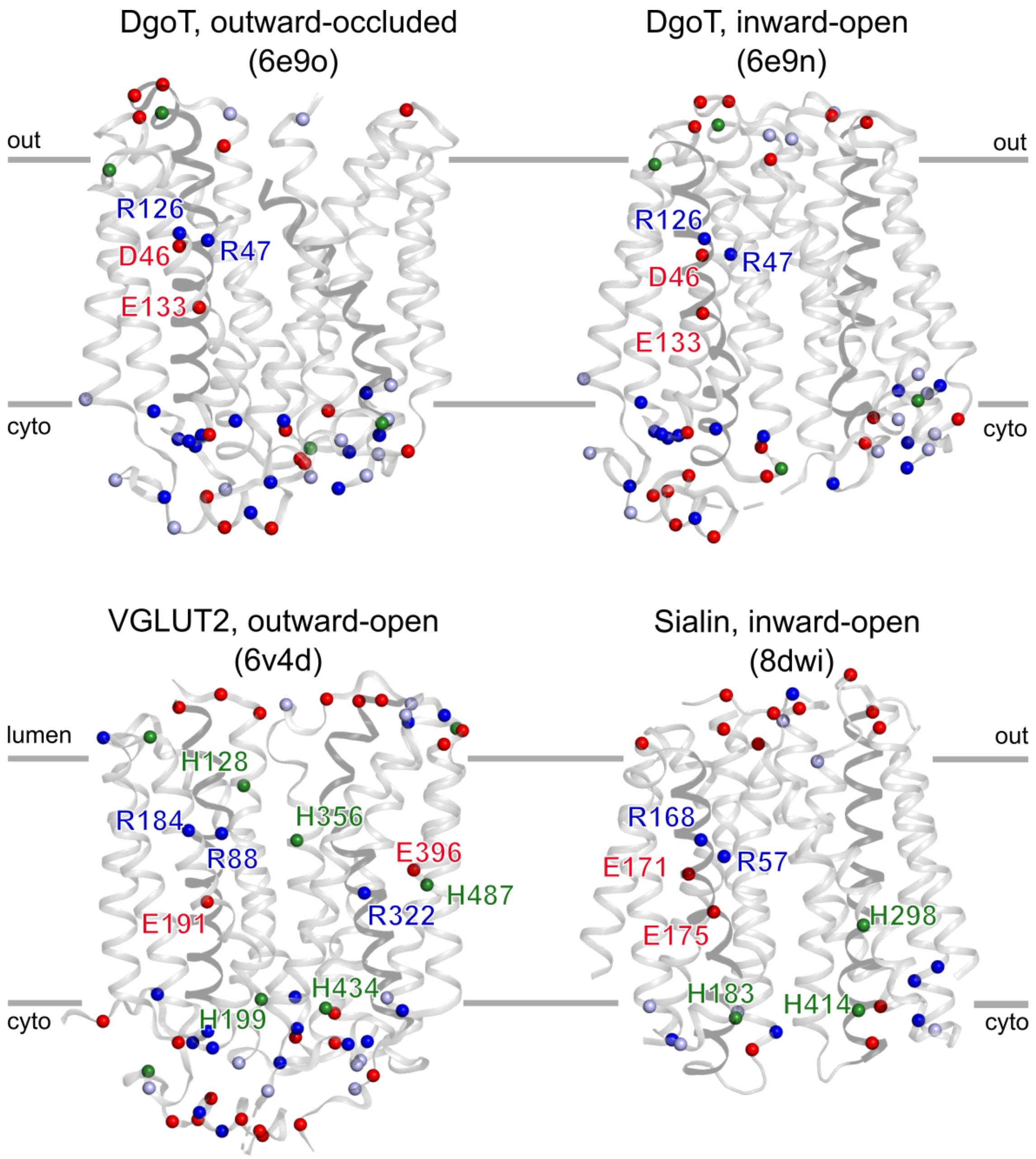


**Supplementary Fig. 1. Structural comparison of SLC17 members DgoT, VGLUT2 and sialin.** PDB codes of the experimentally determined corresponding structures are shown between parentheses. The approximate location of the membrane is indicated with gray lines. Spheres represent the Cα position of Asp and Glu (red), His (green), Arg (dark blue) and Lys (light blue). Charged and titratable residues within the membrane are labeled.
