## Supplementary Fig. 2 for "Transport mechanism of DgoT, a bacterial homolog of SLC17 organic anion transporters"

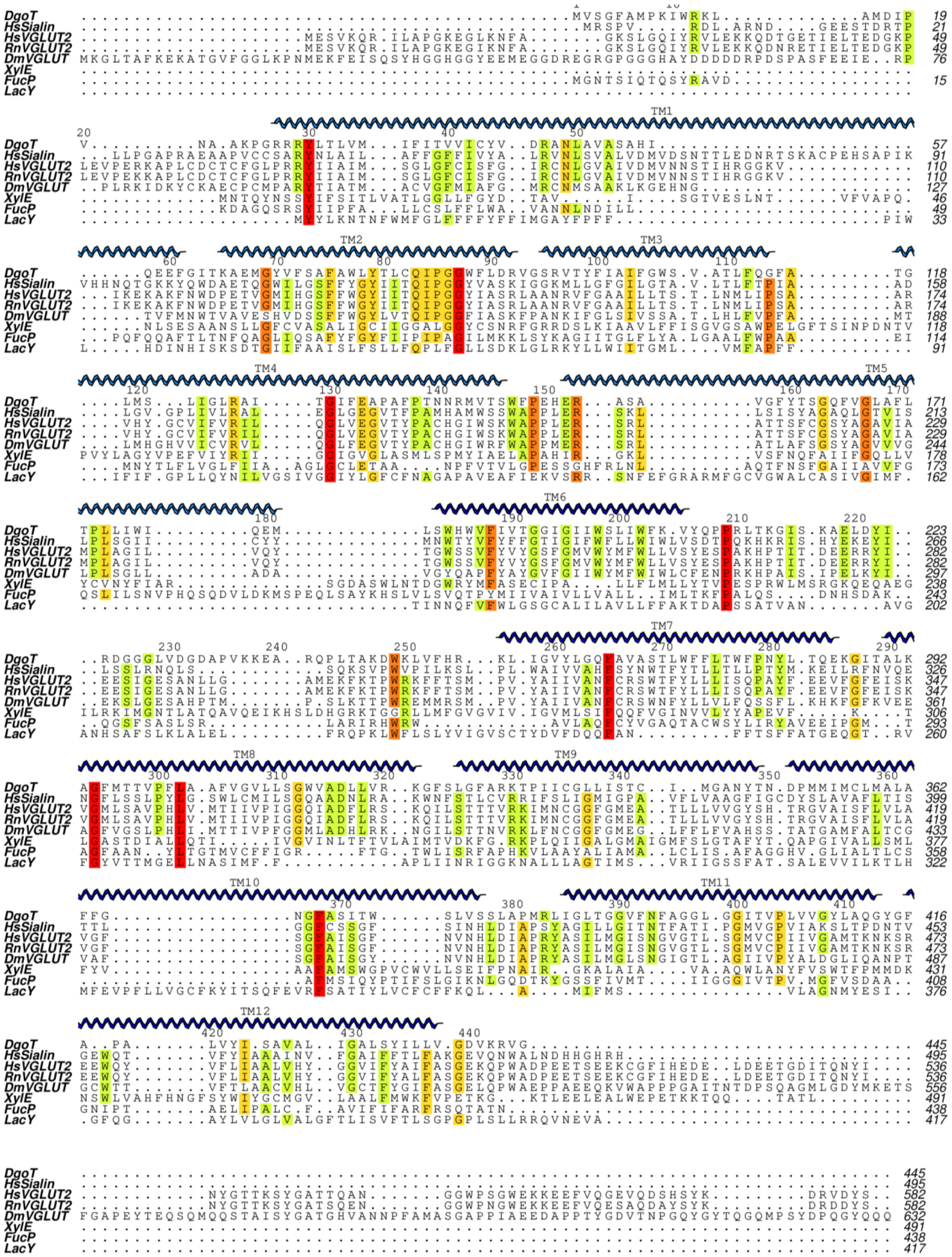


**Supplementary Fig. 2. Sequence alignment of DgoT and related MFS transporters.**

Following protein sequences were used for analysis: D-galactonate transporter from

*Escherichia coli* (DgoT, AKK15832.2), sialin from *Homo sapiens* (HsSialin, CAB62540.1), vesicular glutamate transporter 2 from *Homo sapiens* (HsVGLUT2, NP_065079.1), vesicular glutamate transporter 2 from *Rattus norvegicus* (RnVGLUT2, NP_445879.1), vesicular glutamate transporter from Drosophila melanogaster (DmVGLUT2, AAF51256.2), xylose transporter from *Escherichia coli* (XylE, CAD6020582.1), fucose transporter from *Escherichia coli* (FucP, WP_000528603.1), lactose permease from *Escherichia coli* (LacY, WP_000291549.1).
