## Supplementary Fig. 3 for "Transport mechanism of DgoT, a bacterial homolog of SLC17 organic anion transporters"

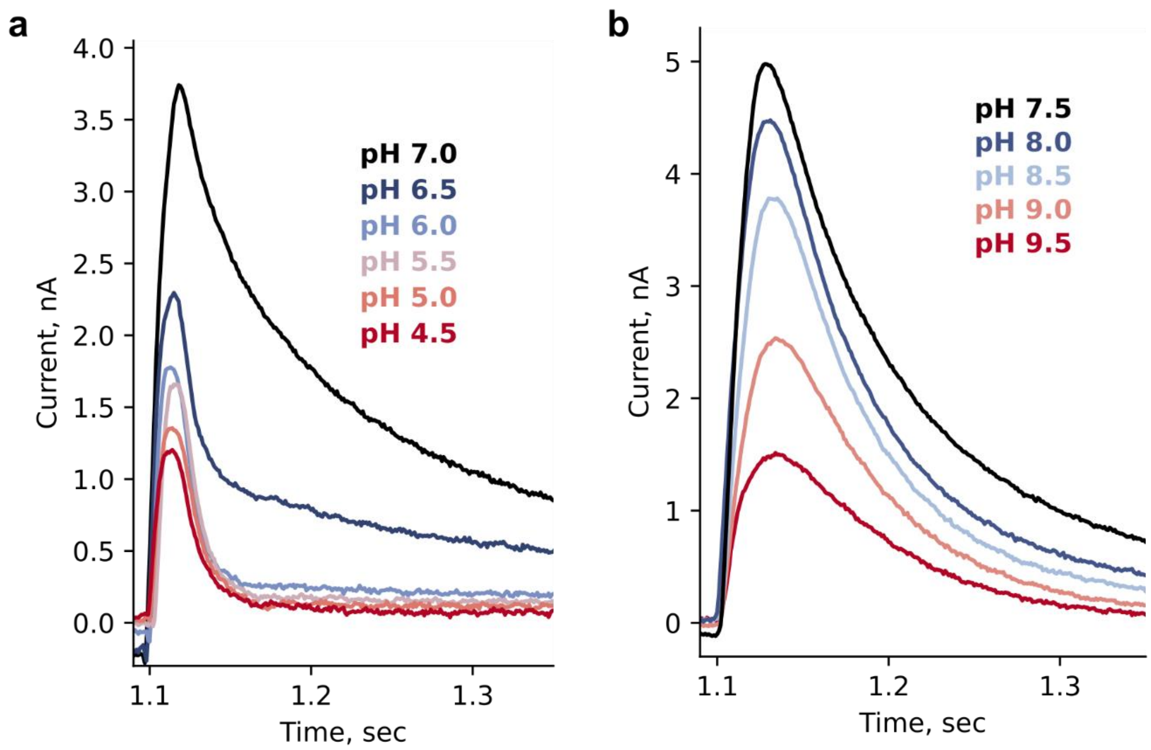


**Supplementary Fig. 3. pH dependence of WT DgoT peak currents measured by SSME upon application of 10 mM D-galactonate concentration jump.** Transient currents obtained using the low time resolution set up (3 mm sensors)
