## Supplementary Fig. 4 for "Transport mechanism of DgoT, a bacterial homolog of SLC17 organic anion transporters"

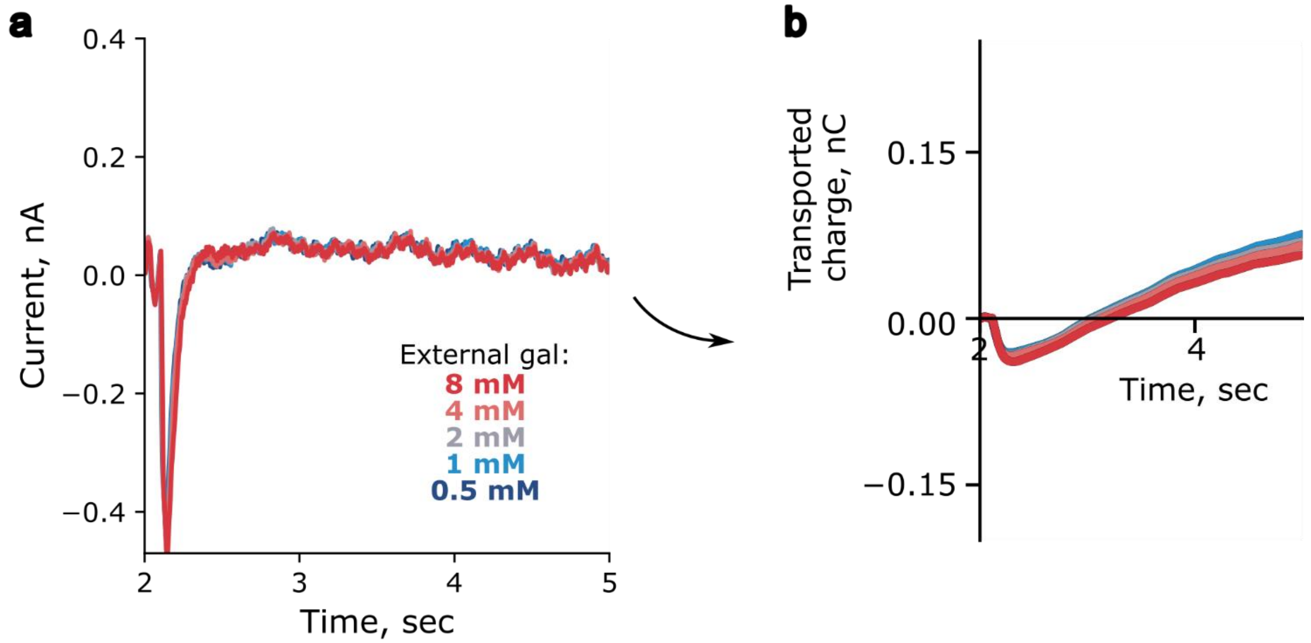


**Supplementary Fig. 4. Current traces recorded with empty liposomes used as a negative control. a** Raw currents recorded with different external solutions. **b** Time dependence of transported charge obtained by integration of current traces.
