## Supplementary Fig. 5 for "Transport mechanism of DgoT, a bacterial homolog of SLC17 organic anion transporters"

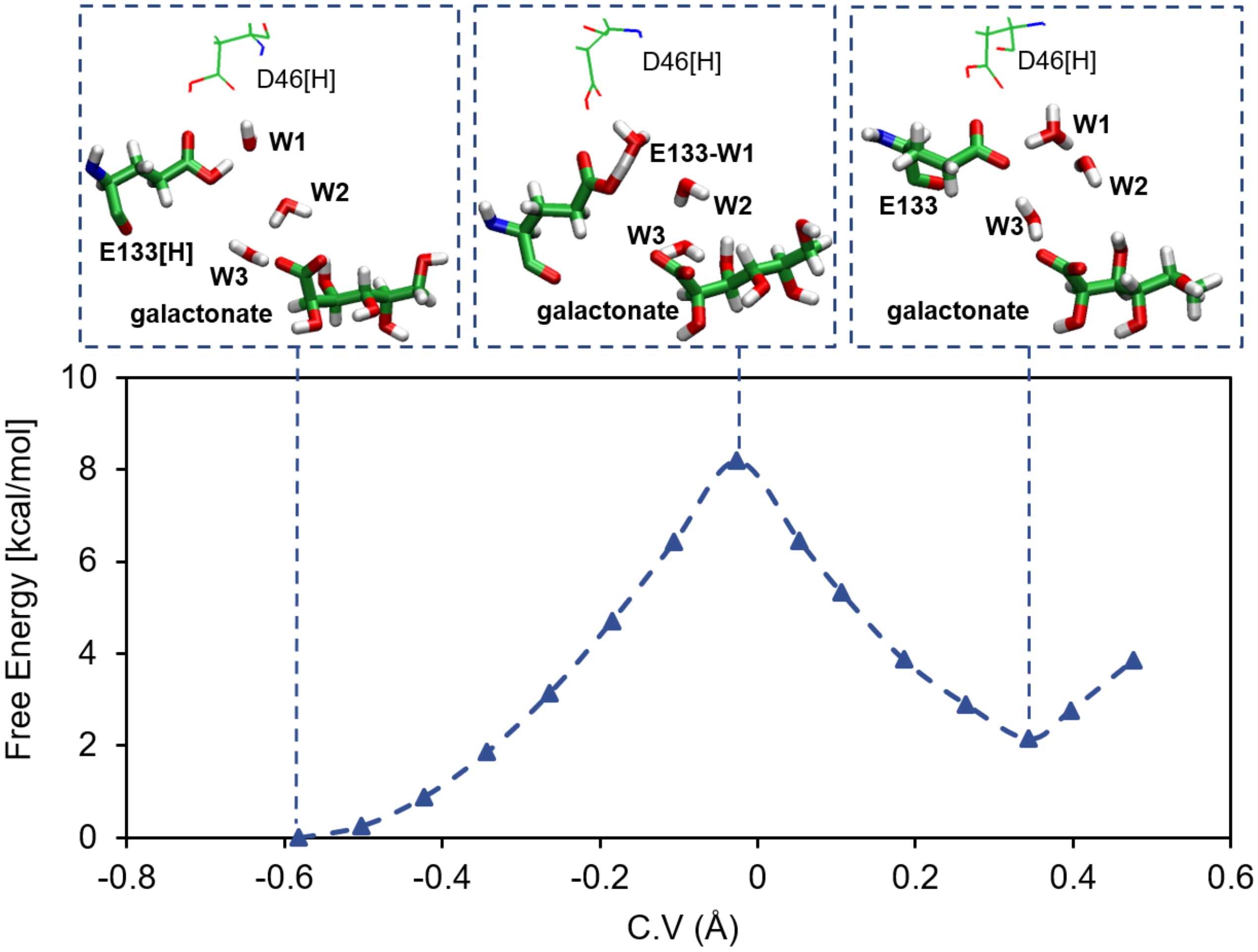


**Supplementary Fig. 5. Free energy profile for the proton transfer between E133 and the adjacent water molecule (W1) in the *close*-D46 configuration, computed at the QM(BLYP)/MM level.** Error bars are omitted since they are smaller than marker size. The insets show representative starting, transition state, and final configurations. QM residues are shown in sticks, while MM residues are shown as lines.
