## Supplementary Fig. 6 for "Transport mechanism of DgoT, a bacterial homolog of SLC17 organic anion transporters"

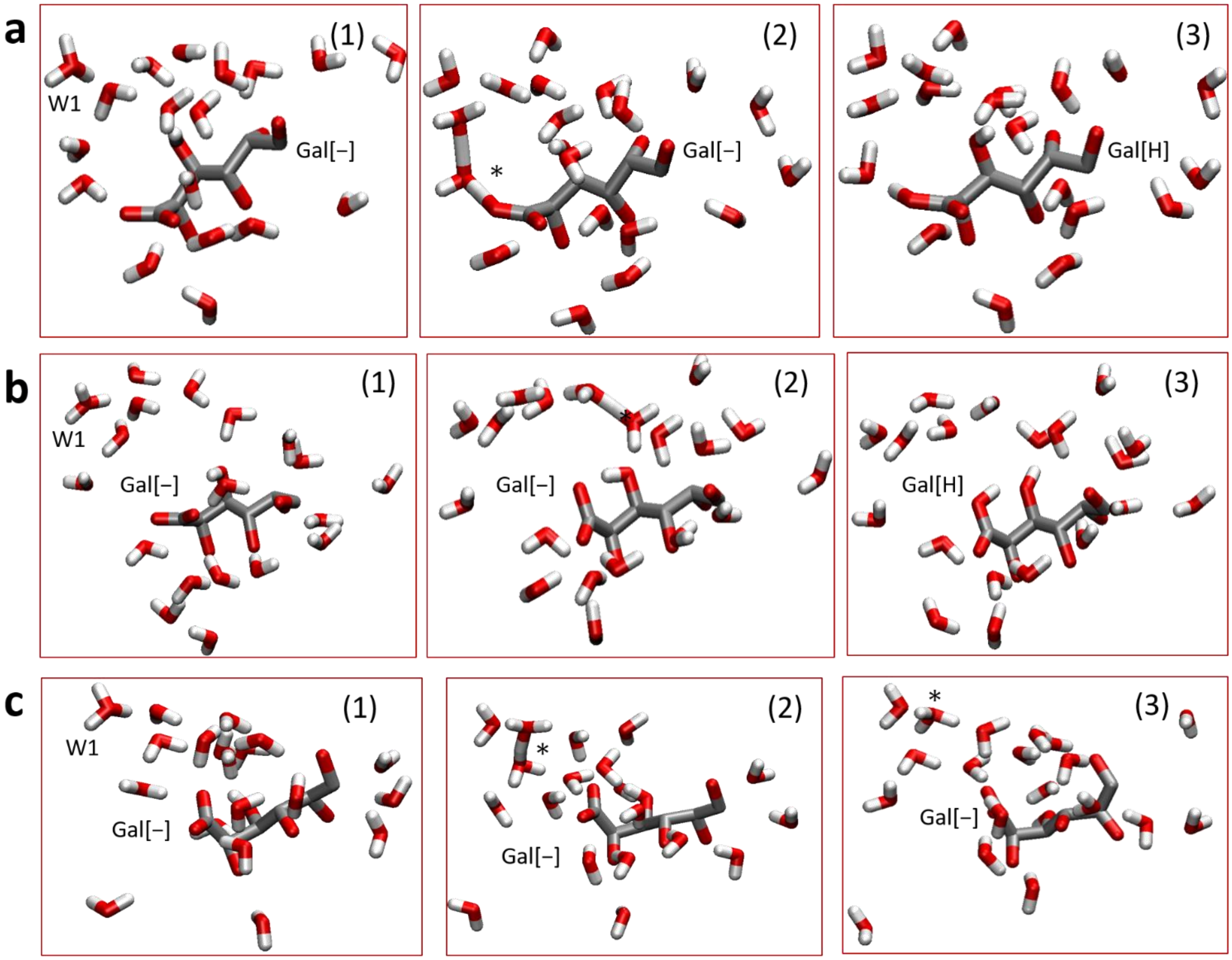


**Supplementary Fig. 6. Representative structures of the QM/MM MD simulations.**

**a** Snapshots of trajectory started from the "snap_4" configuration (see Supplementary Table 2), showing the direct proton transfer from the hydronium ion to galactonate (Gal[H]). Initially the excess proton is located on the first water (W1) and migrates through a 4-water wire until reaching the carboxyl group of galactonate. **b** Snapshots of trajectory started from the "snap_6" configuration, depicting the proton transfer from hydronium ion to galactonate through a 6-water wire and the β-hydroxyl group of galactonate. **c** Snapshots of trajectory started from the "snap_5" configuration, depicting the stablization of the excess protons along the water wire. The asterisk indicates the position of the excess proton in the intermediate structures.
