## Supplementary Fig. 7 for "Transport mechanism of DgoT, a bacterial homolog of SLC17 organic anion transporters"

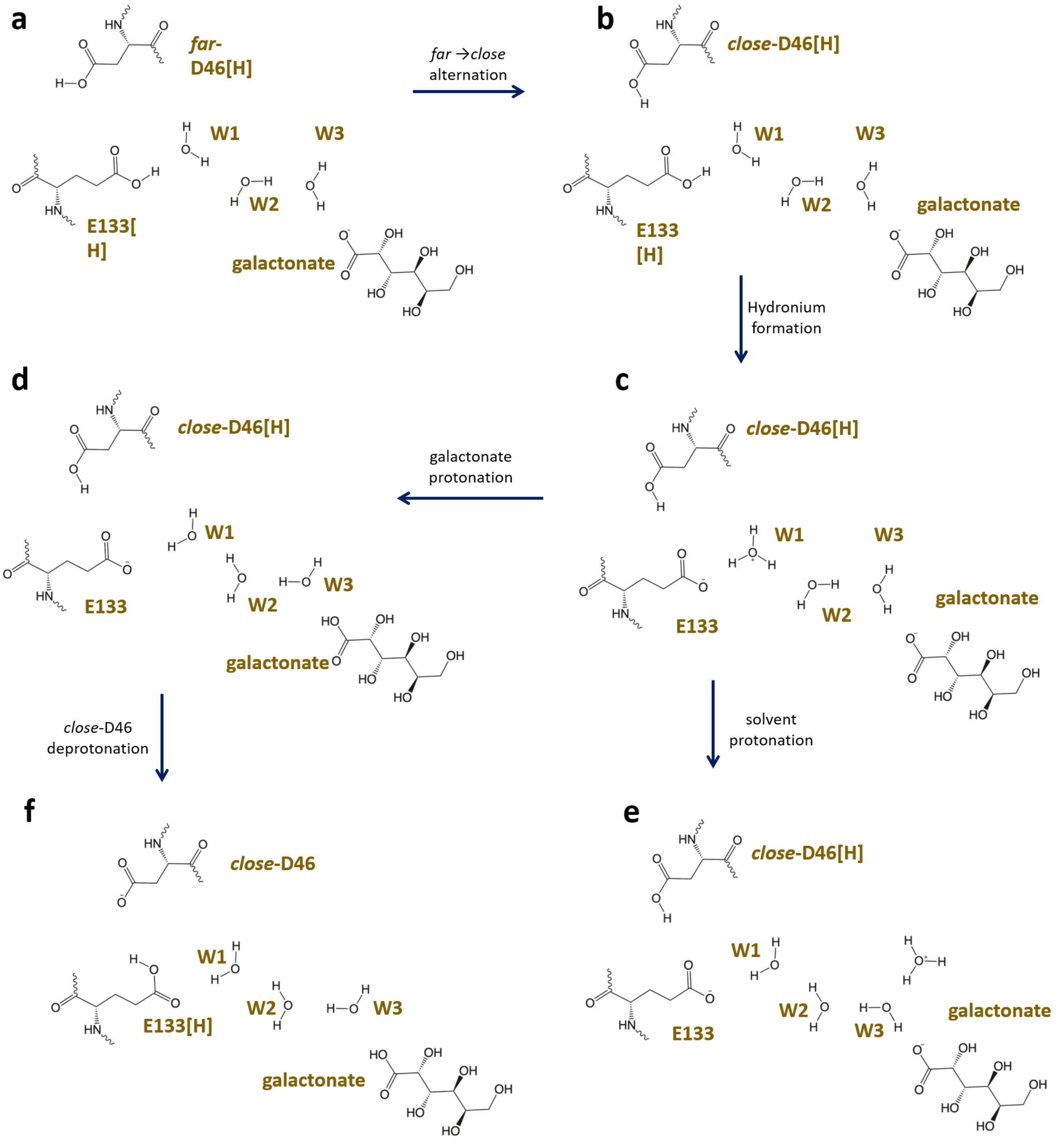


**Supplementary Fig. 7. Schematic Illustration of the proton release in DgoT**. **a** The initial structure featuring protonation of both E133 and D46. **b** Conformational transition of D46[H] from a distant to a close state (*far*→*close*), leading to the formation of a proton pathway between D46 and E133. c Initial proton transfer from E133 to the nearby water molecule (W1), resulting in the formation of a hydronium ion. This excess proton can subsequently be directed either towards the substrate galactonate (**d**) or stabilized within the water network (**e**)**. f** Deprotonation of D46 facilitated by the now deprotonated E133.
