## Supplementary Fig. 8 for "Transport mechanism of DgoT, a bacterial homolog of SLC17 organic anion transporters"

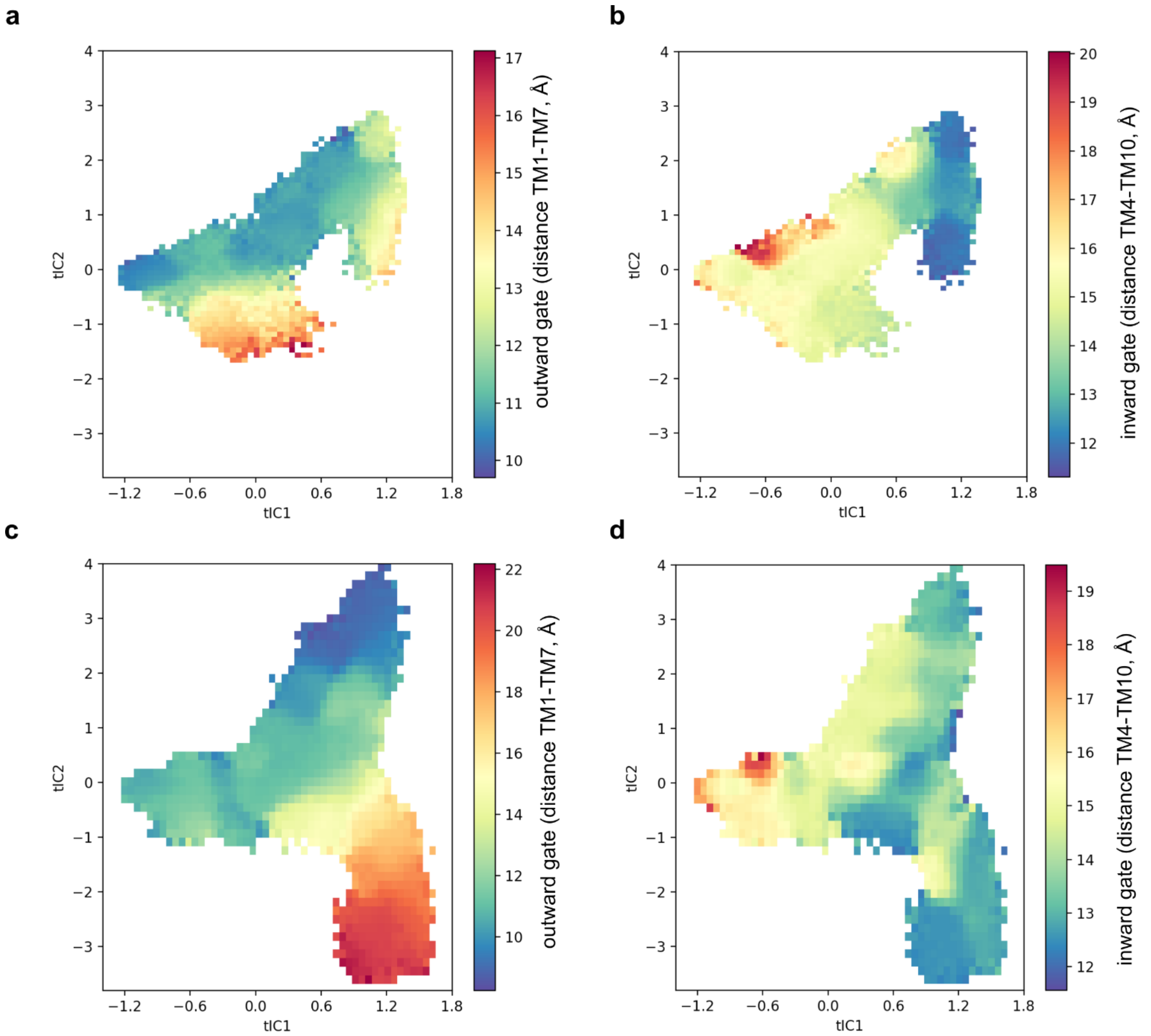


**Supplementary Fig. 8. Correlation between tICA eigenvectors and distances between gating helices. a** and **b** show data for substrate-bound system, **c** and **d** – for *apo* system.
