## Supplementary Fig. 9 for "Transport mechanism of DgoT, a bacterial homolog of SLC17 organic anion transporters"

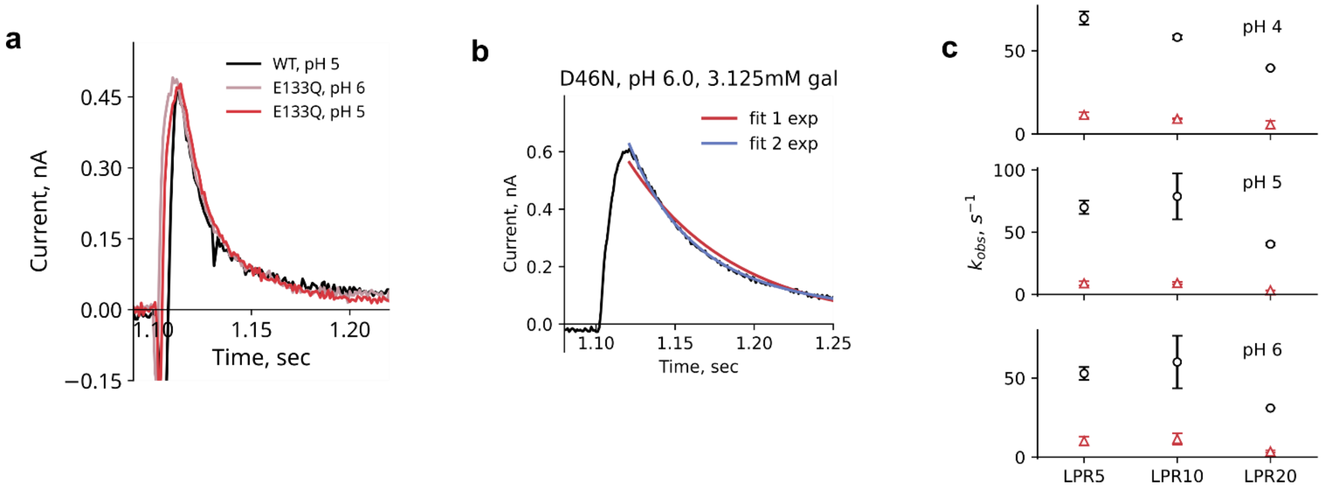


**Supplementary Fig. 9. Pre-steady-state currents recorded with neutralizing mutants DgoT. a** Comparison of representative pre-steady state current recorded with WT and E133Q DgoT at pH 5 or pH 6. **b** Representative D46N DgoT currents with fits to mono- (red line) or biexponential (blue line) functions. **c** Comparison of decay time constants obtained with biexponential fit of currents recorded with D46N DgoT reconstituted in liposomes at different LPR.
