## Supplementary figures and images for "Transport mechanism of DgoT, a bacterial homolog of SLC17 organic anion transporters"

### Supplementary Fig. 10

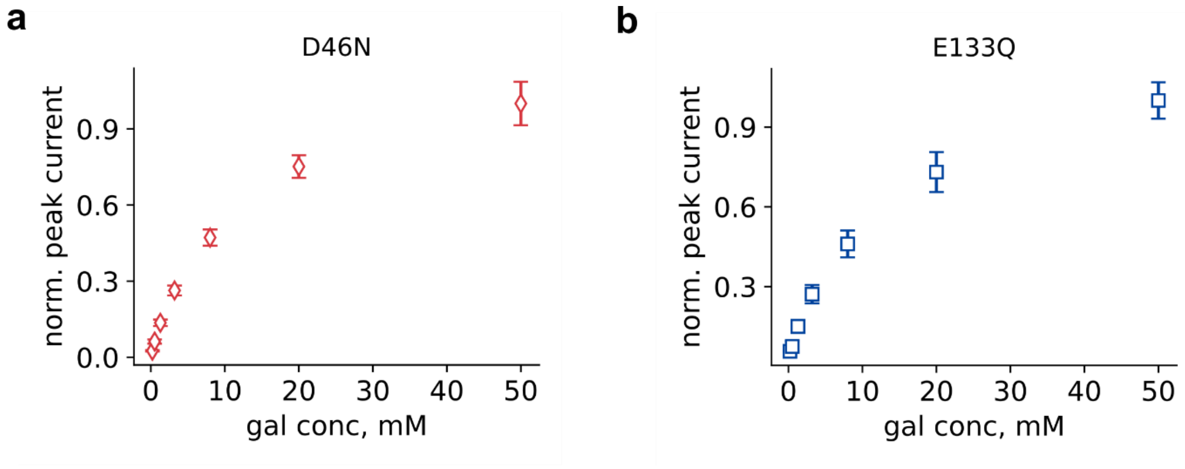


**Supplementary Fig. 10. Normalized peak currents recorded with D46N (a) and E133Q (b) mutants.**
