## Supplementary Fig. 11 for "Transport mechanism of DgoT, a bacterial homolog of SLC17 organic anion transporters"

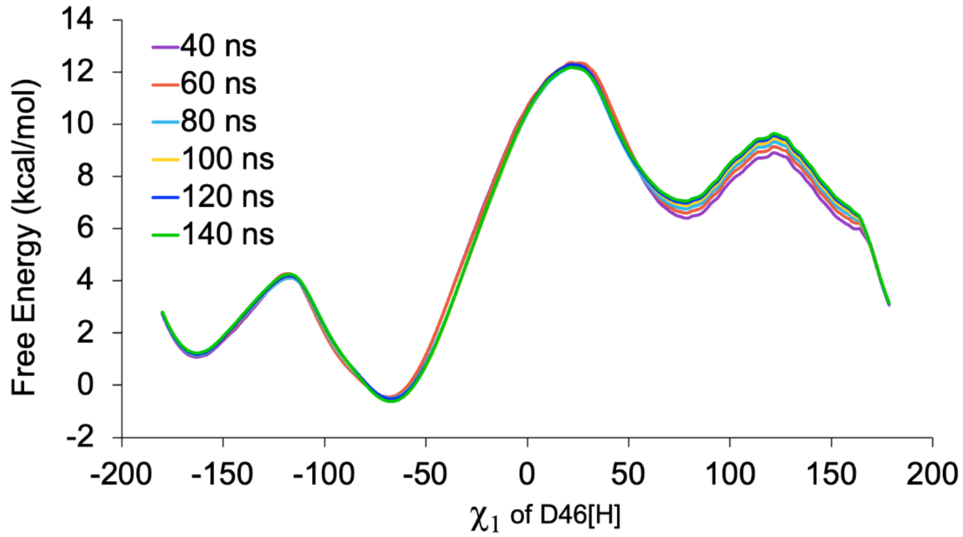


**Supplementary Fig. 11.** Convergence of the free energy profile in classical MTD simulations on the conformational alternation of the protonated D46 as a function of the $\chi_1$ dihedral.
