## Supplementary Fig. 12 for "Transport mechanism of DgoT, a bacterial homolog of SLC17 organic anion transporters"

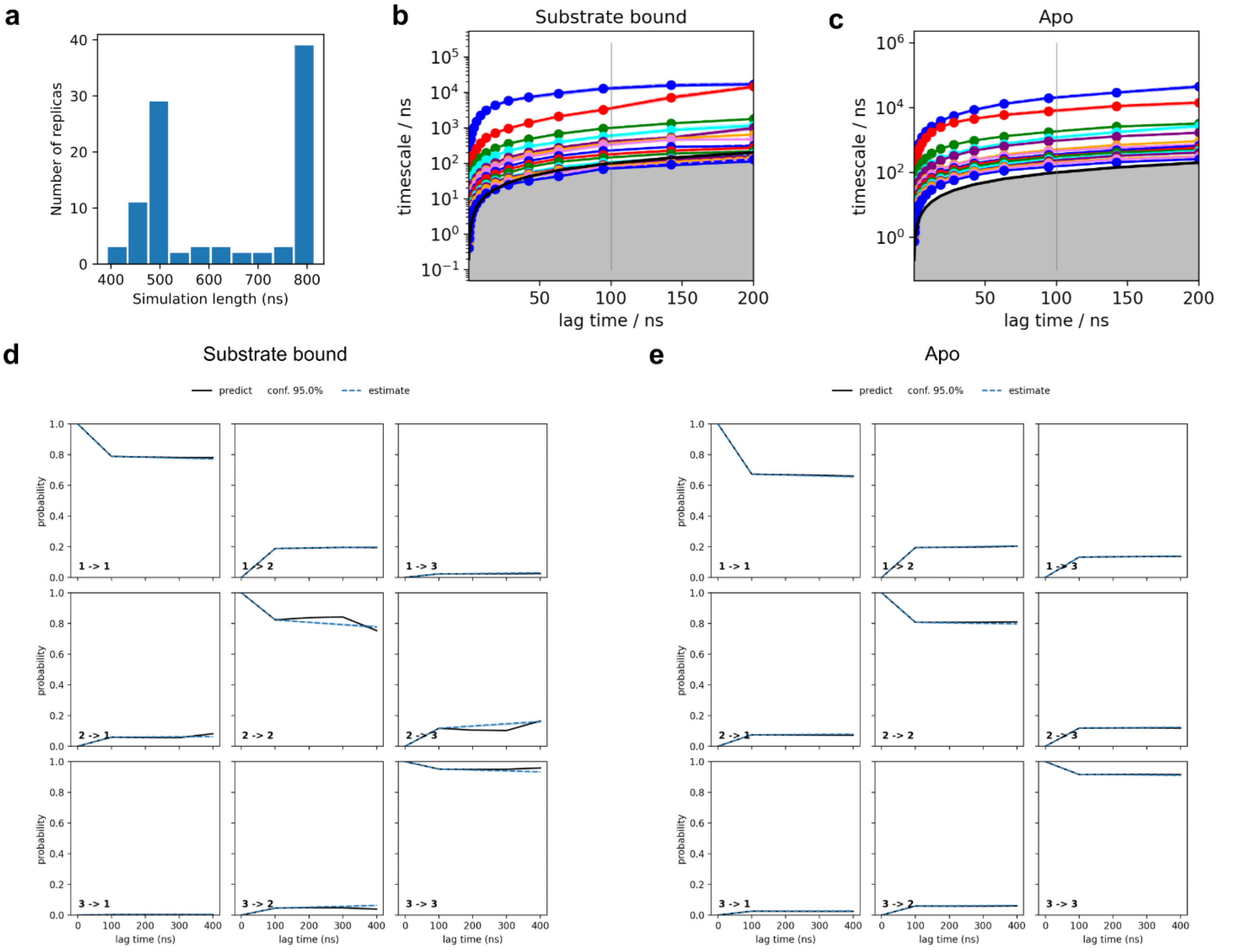


**Supplementary Fig. 12. Estimation and validation of Markov state modeling of DgoT. a** Histogram of lengths of trajectories used for MSM construction. **b, c** Implied timescales plot for systems with substrate bound (b) or *apo* (c) DgoT. Vertical line indicates lag time value chosen for MSM construction. **d, e** Chapman-Kolmogorov test for systems with substrate bound (**d**) and *apo* (**e**) DgoT.
