## Supplementary Table 1 for "Transport mechanism of DgoT, a bacterial homolog of SLC17 organic anion transporters"

| Protonation state: deprotonated (-) or protonated (H) | | | Total number of replica (500 ns long each) | Number of replica where substrate release was observed |
|---|---|---|---|---|
| D46 | E133 | galactonate | | |
| - | - | - | 4 | 2 |
| - | H | - | 10 | 3 |
| H | - | - | 10 | 0 |
| H | H | - | 5 | 0 |
| - | H | H | 4 | 2 |
| H | - | H | 4 | 0 |

**Supplementary Table 1. Substrate release in unbiased MD simulations with inward-facing DgoT with galactonate bound**
