## Supplementary Table 2 for "Transport mechanism of DgoT, a bacterial homolog of SLC17 organic anion transporters"

| Starting configuration | Simulation time (ps) | Proton transfer to galactonate | Water wires | Tau_W1 (ps) | Tau_W2 (ps) | Tau_$W_{gal}$ (ps) | t_gal (ps) |
|---|---|---|---|---|---|---|---|
| snap_1 | 20 | yes | 4 | ≤ 0.025 | 0.125 | 0.125 | 0.45 |
| snap_2 | 30 | shared | 3 | ≤ 0.050 | 0.825 | ≤ 2.10 | 4.5 |
| snap_3 | 35 | no | 3 | ≤ 0.025 | 1.425 | 0.175 | -- |
| snap_4 | 25 | yes | 4 | ≤ 0.050 | 0.275 | ≤ 0.025 | 0.55 |
| snap_5 | 40 | no | 3 | ≤ 0.025 | 40.0 | -- | -- |
| snap_6 | 35 | yes | 6 | ≤ 0.020 | 0.150 | ≤ 0.025 | 14.42 |
| snap_7 | 35 | no | 5 | ≤ 0.020 | 1.025 | 15.750 | -- |

**Supplementary Table 2.** Statistical information pertaining to the proton transfer dynamics simulations for seven representative structures with a hydrated excess proton (snaps 1-7; see Methods). The "Water wire" column denotes the number of water molecules, including the hydronium ion, connecting E133 and galactonate. "Tau_W1" and "Tau_W2" represent the time (in ps) that the proton spends on the initial hydronium ion and its closer neighbor water molecule, respectively, whereas "Tau_$W_{Gal}$" corresponds to the time that the water molecule adjacent to the carboxyl group of galactonate remains as hydronium ion. "t_gal" (in ps) indicates the total time required for the galactonate to undergo protonation.
